# Identifying regions in prefrontal cortex related to working memory improvement: a novel meta-analytic method using electric field modeling

**DOI:** 10.1101/2021.03.11.435002

**Authors:** Miles Wischnewski, Kathleen E. Mantell, Alexander Opitz

**Author notes:** Corresponding author: Miles Wischnewski, PhD. University of Minnesota Department of Biomedical Engineering.

## Abstract

Altering cortical activity using transcranial direct current stimulation (tDCS) has been shown to improve working memory (WM) performance. Due to large inter-experimental variability in the tDCS montage configuration and strength of induced electric fields, results have been mixed. Here, we present a novel meta-analytic method relating behavioral effect sizes to electric field strength to identify brain regions underlying largest tDCS-induced WM improvement. Simulations on 69 studies targeting left prefrontal cortex showed that tDCS electric field strength in lower dorsolateral prefrontal cortex (Brodmann area 45/47) relates most strongly to improved WM performance. This region explained 7.8% of variance, equaling a medium effect. A similar region was identified when correlating WM performance and electric field strength of right prefrontal tDCS studies (n = 18). Maximum electric field strength of five previously used tDCS configurations were outside of this location. We thus propose a new tDCS montage which maximizes the tDCS electric field strength in that brain region. Our findings can benefit future tDCS studies that aim to affect WM function.

**Highlights:**

- We summarize the effect of 87 tDCS studies on working memory performance
- We introduce a new meta-analytic method correlating tDCS electric fields and performance
- tDCS-induced electric fields in lower DLPFC correlate significantly with improved working memory
- The lower DLPFC was not maximally targeted by most tDCS montages and we provide an optimized montage

## 1. Introduction

Working memory (WM) is an executive function allowing for temporary manipulation of information, which is crucial for a large variety of cognitive processes including language comprehension, decision making, learning, and reasoning (Baddeley, 1992). Accordingly, WM deficits are observed in neuropsychiatric disorders, including major depressive disorder, schizophrenia, and obsessive-compulsive disorder (Gold et al., 2019; Heinzel et al., 2018; Marraziti et al., 2010). Therefore, understanding the neural mechanisms underlying WM is crucial to maintain or restore healthy cognitive functioning.

Over the last two decades neuromodulation using transcranial direct current stimulation (tDCS) has been shown to alter cortical activity and consequently human behavior (Polania et al., 2018). When using tDCS, a weak electric current is applied to the scalp, resulting in a low amplitude electric field being induced in the brain non-invasively. By targeting brain regions corresponding to specific cognitive functions, tDCS has the potential to alter behavioral performance (Kuo et al., 2014). Importantly, modeling studies that have simulated the tDCS electric field distribution highlighted the influence of electrode montage and brain anatomy on the affected brain regions (Miranda et al., 2013; Opitz et al., 2015). Thus, for tDCS to alter performance, a montage needs to be used that adequately targets cortical regions underlying specific behavior.

The effect of tDCS on WM performance has been investigated in a large number of studies. Initial meta-analyses have suggested a potential increase of WM performance after anodal tDCS over prefrontal cortex (PFC) (Brunoni & Vanderhasselt, 2014; Hill et al., 2016; Manusco et al., 2016). However, variability across studies was considerable and overall effect size was small. Consequently, different analyses where results are divided into more detailed categories may render meta-analytic findings non-significant (Horvath et al., 2015).

Inter-experimental variability stems from the choice of different tDCS parameters, such as electrode location, targeted area, current intensity, electrode material, electrode orientation, and conductive agent (Polania et al., 2018). Since the classical meta-analysis method generalizes over different features, it is not well-suited to investigate the broad variety in tDCS approaches related to altering WM performance. Moreover, since the publication of previous meta-analytic results on tDCS-related WM effects, numerous new studies have been published exploring a variety of new electrode montages to optimize the effect of tDCS. For instance, recent years have seen a rise in the use of high-definition montages, in which a small stimulation electrode is surrounded by multiple return electrodes to produce a higher focality compared to standard two-electrode montages (Datta et al., 2009; Villamar et al, 2013).

With experimental variability in tDCS montages, induced electric fields vary considerably between studies (Opitz et al., 2018). This issue is further complicated when specific targeted regions differ across studies. For example, to improve WM performance, several studies opted to place the anode over F3 (according to the 10-10 system), in order to target the dorsolateral PFC (DLPFC). However, other studies have used slightly different targets, such as F5 or F7 to stimulate more ventro-lateral portions of the PFC, corresponding to the inferior frontal cortex (IFC) (Di Rosa et al, 2019, Weintraub-Brevda & Chua, 2019). In addition to the variability in location, stimulation intensity strongly differs across studies. Indeed, no normative value for tDCS intensity exists and current intensities typically vary between 0.5 mA and 2 mA across studies. As such, even if differences in WM tasks and outcome measures are set aside, variability in tDCS parameters alone is likely to be a major contributing factor for heterogeneity in results.

To tackle inter-experimental variability and understand tDCS-induced effects on WM, we introduce a novel meta-analytic method to systematically analyze electric field distributions in relation to behavioral performance. With this method, simulated tDCS-induced electric field strength is locally associated with changes in WM performance, providing an estimate for tDCS efficacy at a particular brain region. In other words, our computational meta-analytic method exploits variability in tDCS montage and intensity to form a map of brain regions for which tDCS most strongly correlates to beneficial effects on WM. In order to perform our analysis, a classical meta-analysis was first performed to summarize effect sizes of different studies. This gave an indication if anodal tDCS over left and right PFC generally alters WM performance, regardless of montage. Second, electric field simulations of 69 experiments over left PFC and 18 over right PFC were run to map the different affected cortical areas. Subsequently, electric fields and effect sizes on WM performance were locally correlated and compared to a null-hypothesis model to map out regions that are most susceptible to tDCS-induced WM improvement. Finally, we provide an optimized tDCS montage which maximizes the electric field strength in the region that our meta-analysis identified.

## 2. Methods

The guidelines of the Preferred Reporting Items for Systematic reviews and Meta-Analyses (PRISMA) was used to structure the present meta-analysis (Moher et al., 2009).

### 2.1. Eligibility criteria and search strategy

We included studies in the present meta-analytic computational study if they adhered to the following criteria: 1) Published in a peer-reviewed journal in the English language, with full-text availability. 2) Experimental design included a randomized placebo-(sham) controlled or baseline-controlled design, 3) effect size data was reported or could be calculated from mean and standard deviation or standard error of mean (SEM), presented in the results section, figures, tables, or supplementary material. 4) Reported data was collected from healthy adult participants. 5) Effects of single session tDCS were reported. From multi-session tDCS studies, only first session results were included (Lally et al., 2013; Looi et al., 2016; Ke et al., 2019; Martin et al., 2013; Richmond et al., 2014; Wang et al., 2019).

We conducted a literature search within databases PubMed and Web of Science in a period between January 2000 and October 2020. The search terms ‘tDCS’ + ‘working memory’ and ‘transcranial direct current stimulation’ + ‘working memory’ were used and the reference sections of research and review articles were examined for additional articles. After removing duplicates, 281 articles were inspected for eligibility, finally yielding 68 selected articles (Supplementary Figure 1A). From these we extracted 69 effect sizes related to left PFC anodal tDCS and 18 effect sizes related to right PFC anodal tDCS (Table 1 and 2).

**Table 1.**
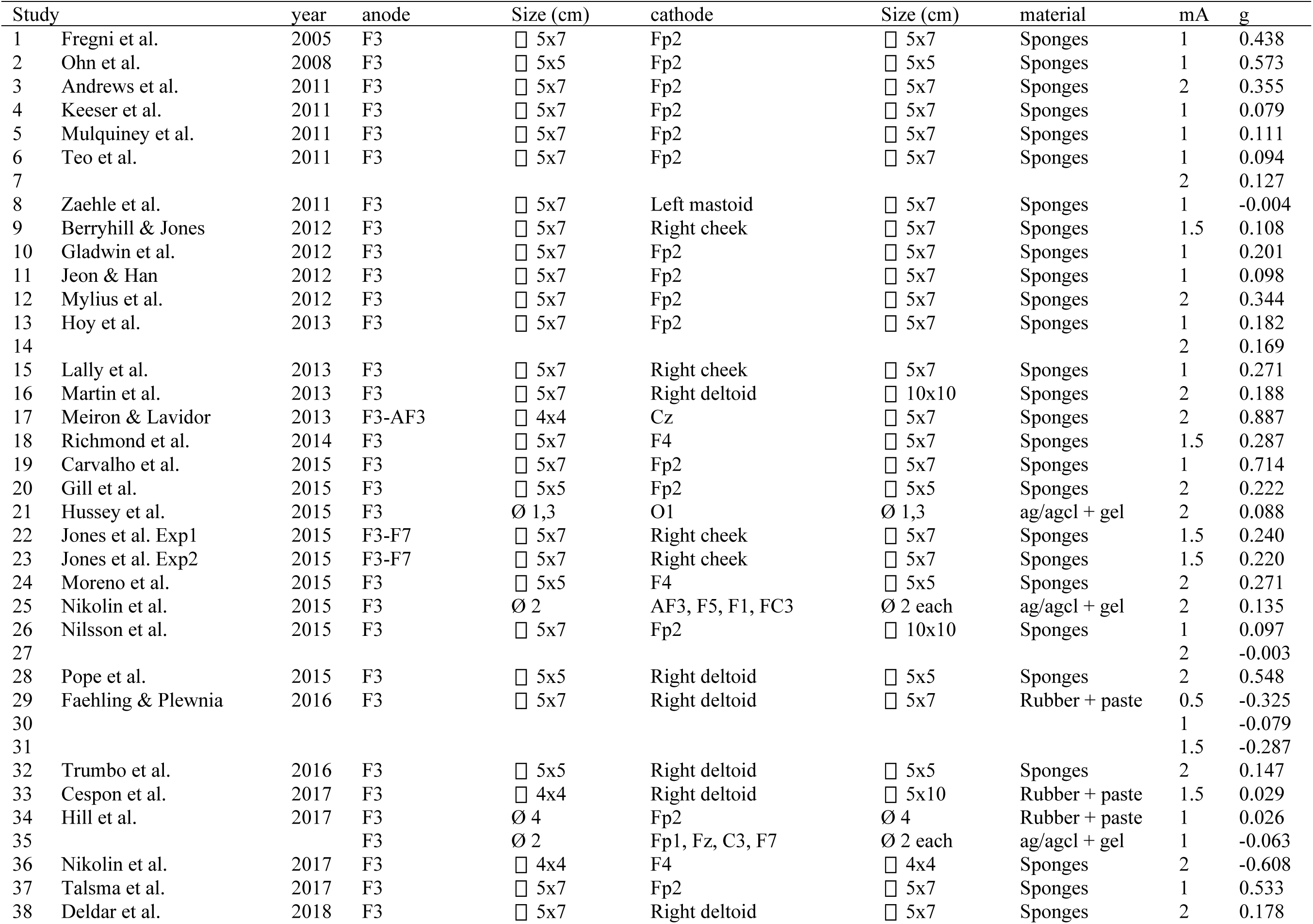
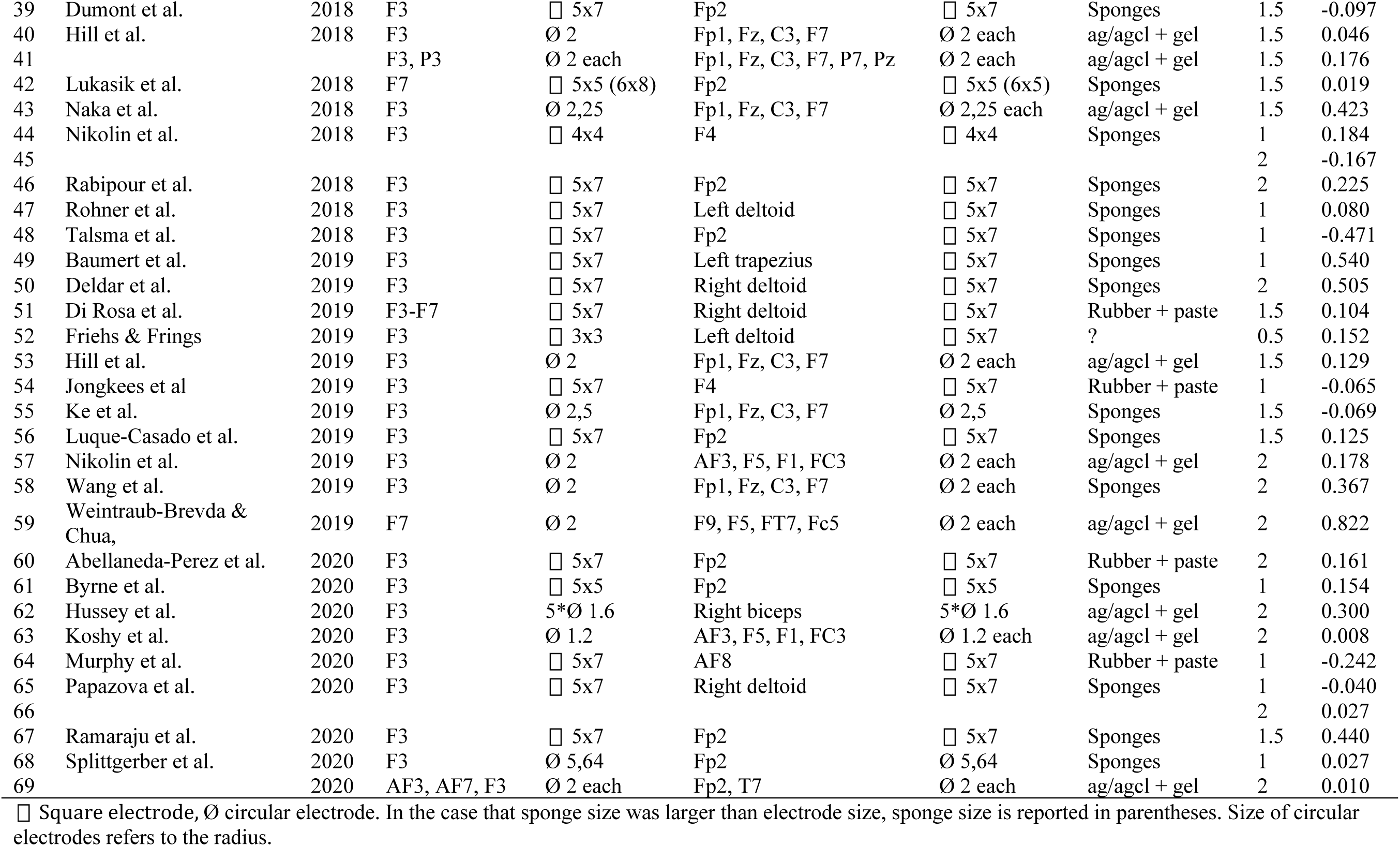
Study overview left PFC anodal tDCS

### 2.2. Outcome variables for effect size calculation

In the present analysis WM was treated as a single cognitive construct and thus we included data from different WM tasks. The studies we investigated used one or more of the following tasks: N-back task, Sternberg task, Corsi block tapping task, paced auditory serial addition and/or subtraction task, digit span, change detection task, internal shift task, delayed WM task, and other custom WM tasks. Different cognitive modalities, such as spatial, visual, auditory, and verbal WM task were included. Also, tasks of various difficulty were included, such as 2-back as well as 4-back tasks (Supplementary Table 1 and 2). However, note that specifically for the n-back task, 0-back and 1-back trials were excluded from analysis here as they are typically employed as a control task and task performance is at ceiling level (Gill et al., 2015). Furthermore, WM tasks with experimental manipulations such as inducing stress (Ankri et al., 2020; Bogdanov & Schwabe, 2016), emotional load (Faehling & Plewnia, 2016; Weintraub-Brevda & Chua, 2019; Wolkenstein et al., 2014), or interference (Gladwin et al., 2012) were included if WM remained the primary tested construct of interest. Outcome measures included accuracy, reaction time, maximum achieved n, sensitivity, forward span, and backward span.

Combining various WM tasks and outcome measures lowers specificity in detecting tDCS-related effects on specific features (e.g. spatial vs verbal WM, or n-back vs Sternberg performance, or accuracy vs reaction time). However, comparison of different WM aspects lowers the amount of observations per category and consequently strongly reduces statistical power. Therefore, we want to emphasize that investigation of tDCS parameters is the goal of the present study and analysis of WM sub-components is beyond the scope of the present study.

### 2.3. Effect size extraction and statistical analysis

We used anodal tDCS-induced WM task performance change compared to a control condition for effect size calculation. Comparison to a control condition was defined as I) the difference between anodal and sham tDCS; or II) if a baseline measurement was present, the difference between baseline corrected anodal (posttest – pretest) tDCS and baseline corrected sham (posttest-pretest) tDCS. Hedges’ g was used as the effect size measure, which is based on Cohen’s d with an adjustment to account for inflation due to small sample sizes (Hedges & Olkin, 1985). We gathered data for effect size calculation from results sections and tables or estimated them from figures and appendices using WebPlotDigitizer 4.3. software (https://github.com/ankitrohatgi/WebPlotDigitizer). Hedges’ g for all outcome measurements (accuracy, reaction time, digit span, etc.) was derived from reported Cohen’s d values or calculated using means and pooled standard error of mean. Subsequently, for each study we pooled effect sizes from different outcome measures, such that a single Hedges’ g value was associated with each tDCS montage and intensity configuration. That is, studies using a single tDCS setup are reflected by a single Hedges’ g value and studies using multiple tDCS configurations are associated with as many effect size values (e.g. Faehling & Plewnia, 2016; Hill et al., 2017; Hoy et al., 2013). A positive Hedges’ g reflects a tDCS-induced increase WM performance, whereas negative g indicates a tDCS-induced decrease of WM performance compared to (baseline-corrected) sham values.

We performed classical meta-analyses using MetaWin 2.1. (Rosenberg et al., 2000) and JASP 0.14. For Hedges’ g from left PFC tDCS (n = 69) and right PFC tDCS (n = 18) a random effects model resulted in the cumulative effect size (Ḡ) and 95% confidence intervals. From this we calculated the corresponding Z-statistic and p-value to investigate whether Ḡ differed significantly from zero. For comparison, effect sizes were calculated for five tDCS montage categories targeting the left hemisphere: PFC-supraorbital region (SOR) (n = 28), PFC-Cheek (n = 4), PFC-Shoulder (n = 12), PFC high-definition (HD) (n = 11) and PFC bifrontal (n = 6). Due to limited sample size this analysis was not performed for experiments on right PFC.

Total heterogeneity (Qt) of effect size distribution was tested (Hedges, 1981) and checked for normality using a Kolmogorov–Smirnov test. Furthermore, the Rosenthal method (α < 0.05) was used to calculate a fail-safe number, which reflects the number of null findings that are necessary to render Ḡ non-significant (Rosenthal, 1979). Symmetry between sample size and mean effect size, that is a funnel plot, was used to assess publication bias (Sterne et al., 2011). Additionally, sample size bias and recency bias were explored by correlating number of participants, publication year, and effect size using a non-parametric Spearman rank-order test (ρ)(Begg & Mazumdar, 1994; Schäfer & Schwarz, 2019).

### 2.4. Risk of bias assessment

We performed an assessment of bias risk on the included studies employing the Cochrane Collaboration’s tool for risk of bias in randomized trials (Higgins et al., 2011). The risk of selection bias, performance bias, detection bias, attrition bias, reporting bias, and other biases was judged for each study and classified as high, low, or uncertain. Results are presented in Supplementary Figure 1B.

### 2.5. FEM Modeling

All FEM simulations were run using SimNIBS version 3.2. (Thielscher et al., 2015). We simulated the tDCS electric field distribution for each included study (left PFC n = 69, right PFC n = 18; Supplementary Figure 2). Specific tDCS montage (electrode location, size, shape and orientation), intensity, electrode material (sponge, rubber, Ag/AgCl), as described in each study, was used for simulations. Simulations were performed on an individual head model of a healthy adult male provided by SimNIBS (“Ernie”). Previously established realistic conductivity values of different tissue types were used: σ_skin_ = 0.465 S/m, σ_bone_ = 0.01 S/m, σ_cerebrospinal fluid_ = 1.654 S/m, σ_gray matter_ = 0.275 S/m, and σ_white matter_ = 0.126 S/m (Windhoff et al., 2013). Gray matter volume was extracted for calculation of electric field strength.

Besides simulations for each study, we created an averaged electric field distribution for five tDCS montage categories commonly used by previous studies (PFC-SOR, PFC-Cheek, PFC-Shoulder, PFC HD, PFC bifrontal), as well as for the entire dataset. The robust maximum of electric field strength (E_MAX_) was quantified as the 99.9^th^ percentile of electric field strength and corresponding MNI coordinates were determined. Affected volume was quantified as the volume (in cm^3^ and percentage of total brain volume) corresponding to half of the maximum electric field (Foc_50_). E_95_, E_75_, E_50_, and Foc_75_ are reported in Supplementary Table 2. E_MAX_ and Foc_50_ were compared between montage categories using repeated-measures ANOVAs, followed by Bonferroni-corrected post-hoc t-tests.

### 2.6. Meta-analytic covariance between electric-field distribution and behavioral performance

We introduce a novel method that allows for associating meta-analytic behavioral effect size and tDCS-induced electric field distributions (Figure 1). To this end we locally correlated electric field strength and Hedges g values across all studies at each gray matter tetrahedron (left PFC, n = 69 and right PFC, n = 18). This correlation will be referred to as the performance – electric field correlation (PEC). PEC values above zero reflect a positive association between electric field and hedges g. This indicates the tDCS-induced electric field strength in a particular brain region relates to increased WM performance. A PEC below zero reflects a negative association between electric field and hedges g. This means that a higher tDCS electric field in a particular brain region relates to decreased WM performance. PEC values were calculated for all tetrahedra resulting in a map displaying the relationship between tDCS-induced electric fields and behavioral change.

**Figure 1.**
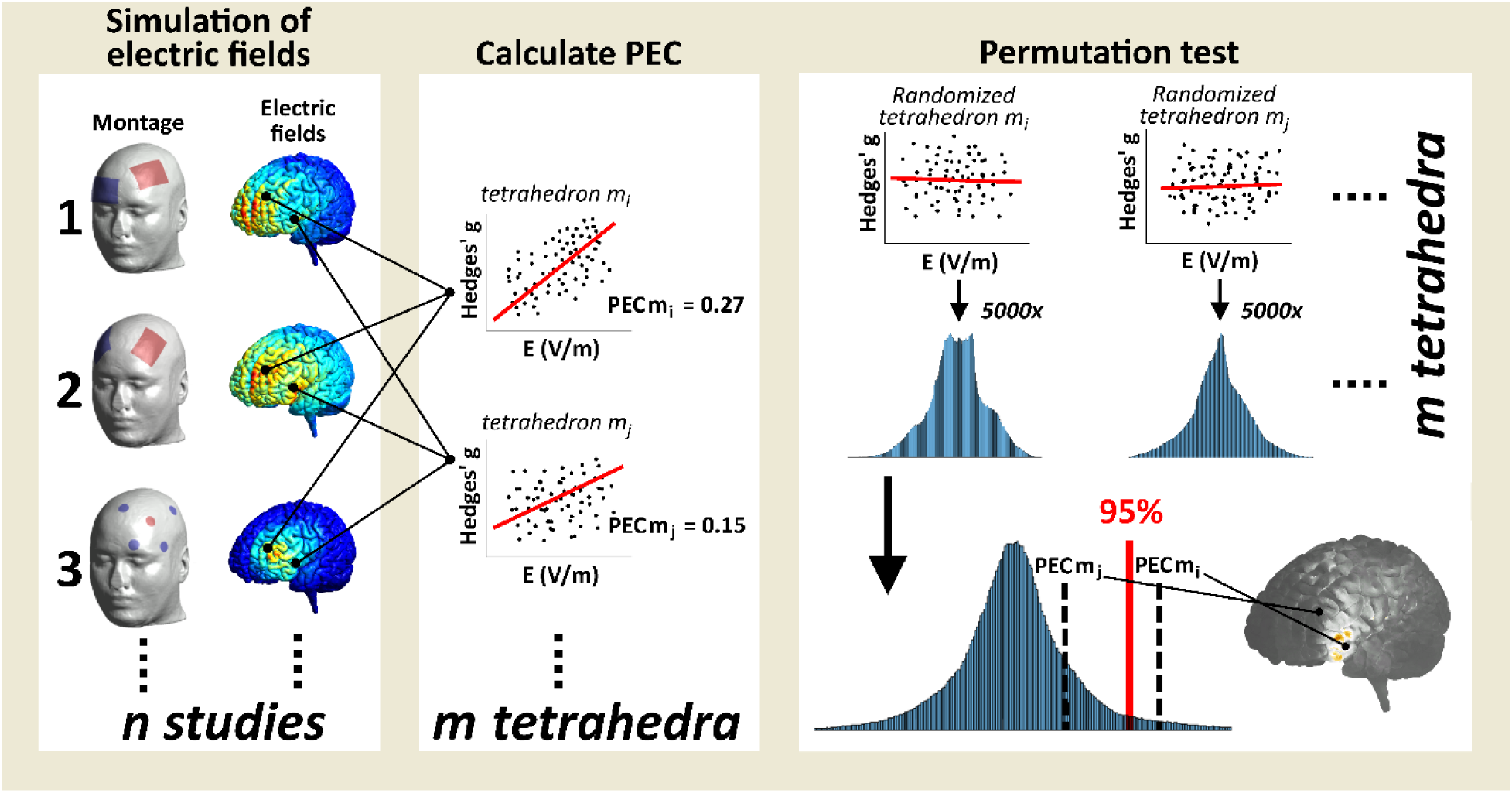
Analysis pipeline for relating electric field distributions to WM performance. First electric fields were simulated for all studies (left PFC n = 69; right PFC n = 18). Subsequently, for each tetrahedron (individual volumetric element in gray matter, m = ∼9.8 x 10^5^) the correlation between electric field strength (V/m) and WM effects size (Hedges’ g) is calculated. This correlation is referred to as the performance – electric field correlation (PEC). To test for significance, a permutation test is performed in which PEC values are compared to a null-hypothesis model. The null-hypothesis model is generated at each tetrahedron by performing 5000 randomized correlations between the shuffled Hedges’ g value and electric field strength. The actual obtained PEC values are compared to the distribution of the null-hypothesis model yielding a (one-sided) t-statistic and corresponding probability (p) value, such the 95^th^ percentile of the distribution corresponds to p = 0.05. Finally, all PEC and inverted log10-transformed p-values are displayed on the gray matter volume in Figure 5 (left PFC) and Figure 7 (right PFC).

After PEC calculation we used a permutation test to statistically analyze PEC values and determine significance compared to a null-hypothesis model (Figure 1). First, the null-hypothesis model was formed by performing 5000 permutations of randomized PEC values at each tetrahedron resulting in an approximate Gaussian distribution. Subsequently, we compared actual obtained PEC values to the null-hypothesis model for each brain region yielding a t-statistic and a corresponding p-value. As we investigated tDCS-induced performance improvement, a one-sided distribution was used, such that the 95% of the Gaussian distribution equals a p-value of 0.05. For presentation purposes, we displayed inverted log_10_ transformed p-values to emphasize low p-values (Figures 5B and Figures 7B).

**Figure 5.**
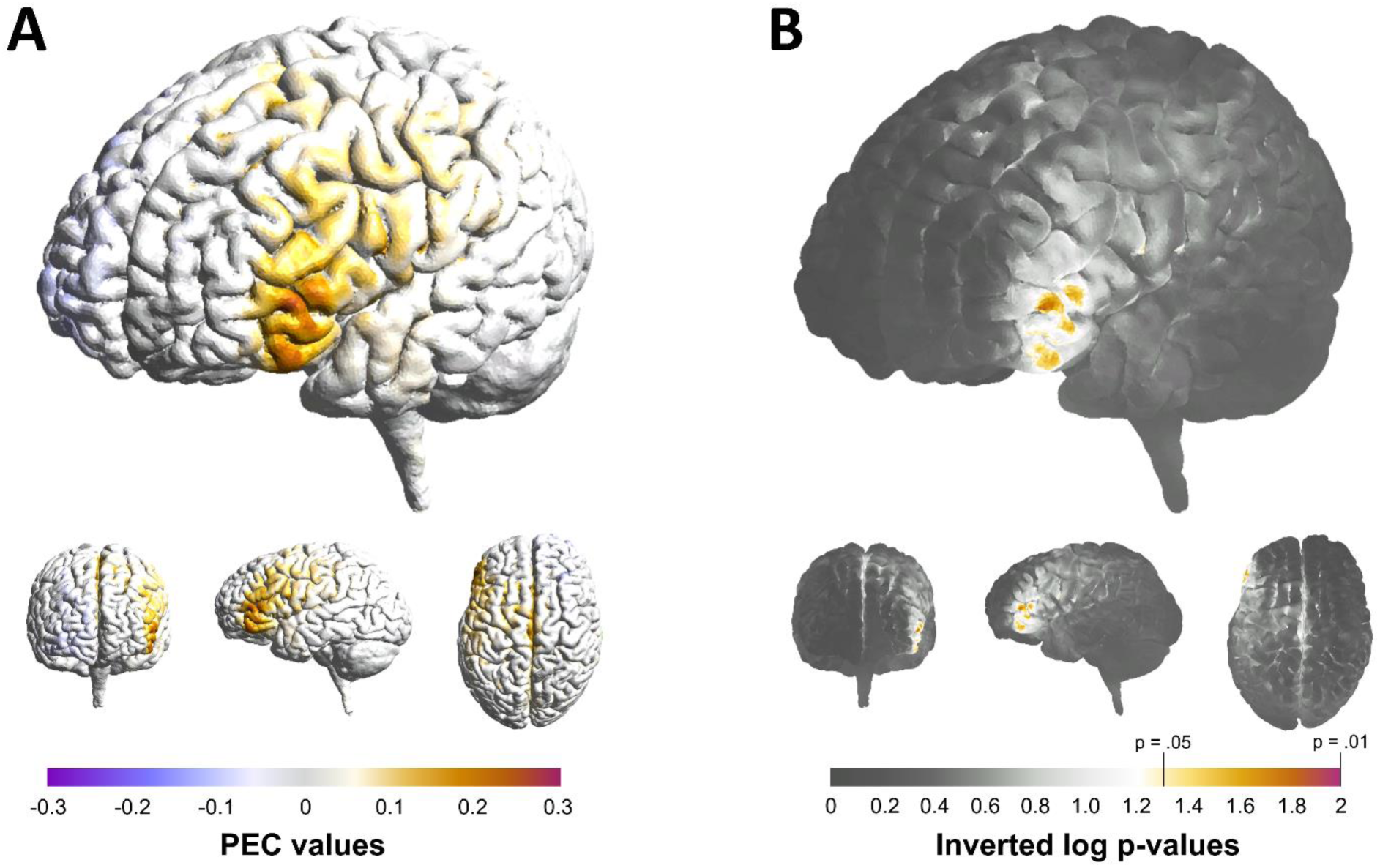
A) PEC values of left PFC tDCS studies representing the relationship between electric field strength and behavioral effect size. That is, higher PEC values indicate that higher electric field strength in a particular brain location relates to higher increased WM performance. B) Inverted Log_10_ p-values associated with the PEC values. Inverted Log-p of 1.31 and 2 correspond to p = 0.05 and p = 0.01 respectively.

**Figure 7.**
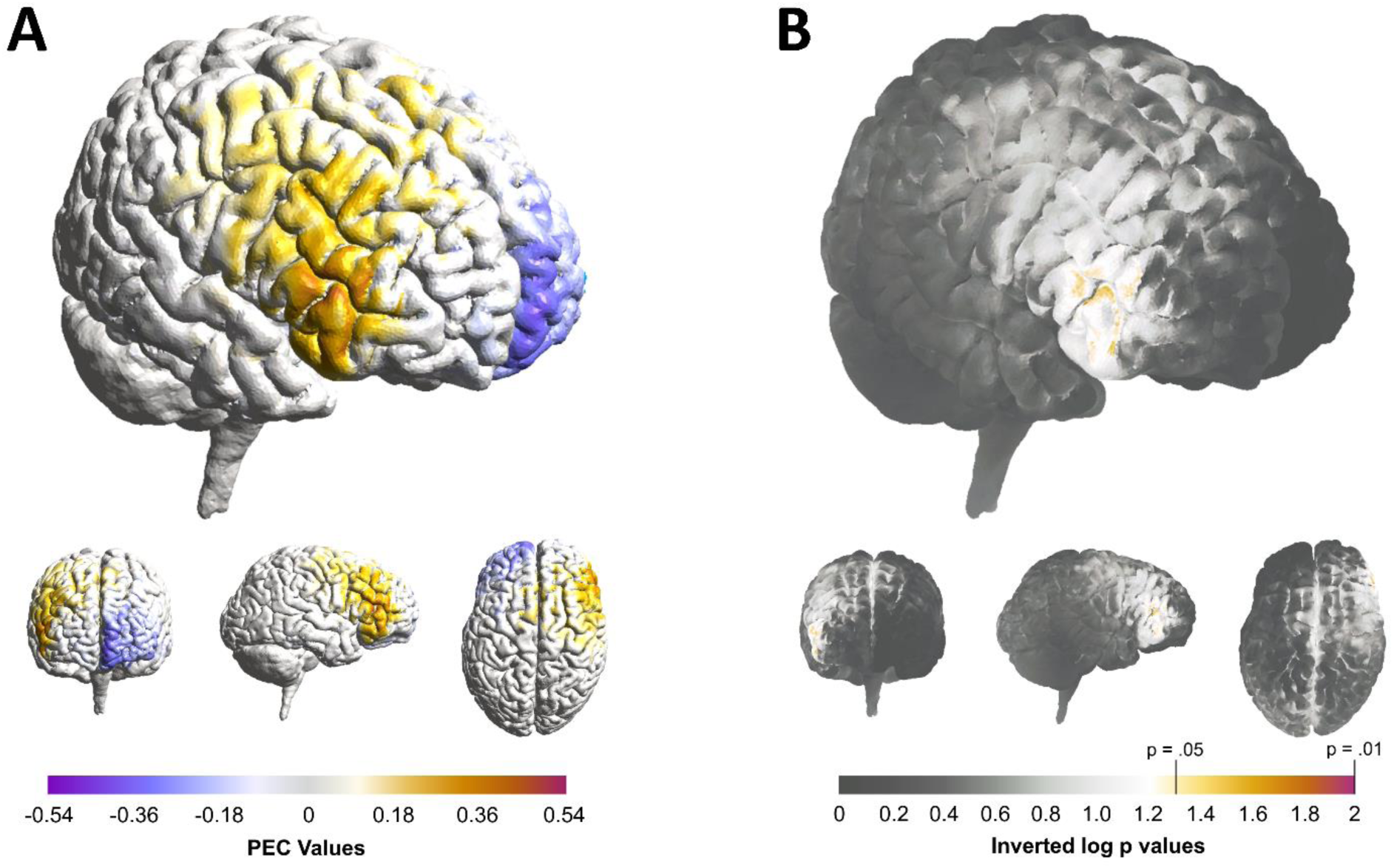
PEC values of right PFC tDCS studies representing the relationship between electric field strength and behavioral effect size. That is, higher PEC values indicate that a higher electric field in a particular location relate to higher increased WM performance. B) Inverted Log_10_ p-values associated with the PEC values. Inverted Log-p of 1.31 and 2 correspond to p = 0.05 and p = 0.01 respectively.

To get an indication of the efficacy of most commonly used tDCS montages, the electric field strength was calculated averaged for all tetrahedra in the region where PEC values are significant. Additionally, the PEC value was determined for each montage category at the location of their respective E_MAX_. Each PEC value was compared to the robust maximum (PEC_MAX_).

### 2.7. tDCS montage optimization

Using the SimNIBS optimization routine (Saturnino et al., 2020), we determined a tDCS montage which maximizes the electric field strength in the left PFC robust maximum PEC (PEC_MAX_) location. The optimization algorithm is performed on the same standard head model (“Ernie”) and 74 possible electrode positions according to the 10-10 EEG system included with SimNIBS. All brain locations in a radius of 10 mm around the PEC_MAX_ location were used as target, with no restraints on electric field direction. The number of stimulation electrodes was set to 5 or fewer, with a maximum intensity of 1 mA per electrode and 2 mA total current. The perpendicular component of the induced electric field was separated at the brain surface level to display inward and outward currents of the optimized tDCS montage.

## 3. Results

### 3.1. Classical meta-analysis for left prefrontal tDCS

We performed a meta-analysis on the effect of left prefrontal tDCS on WM performance to get an indication of general tDCS efficacy irrespective of montage and intensity. A total of 69 effect size values (hedges *g*) of left PFC anodal tDCS studies were collected. Overall, we observed a significant cumulative effect size of Ḡ = 0.147 (95% CI = 0.067 – 0.228), Z = 3.595, p < 0.001, indicating that tDCS has a small beneficial effect on WM performance compared to a control condition (Figure 2A). Total heterogeneity of effect sizes was not significant (Q_T_ = 40.48, p = 0.997) and Kolmogorov–Smirnov test indicated no deviation from normality (D = 0.134, p = 0.156). The fail-safe number indicated that 306 null results would be necessary to render the cumulative effect size non-significant. Effect sizes for tDCS montage categories separately varied between Ḡ = 0.018 to Ḡ = 0.241 (Table 3). Given the smaller sample sizes the compared effectiveness of each montage category is less conclusive, and no montage category appears superior.

**Figure 2.**
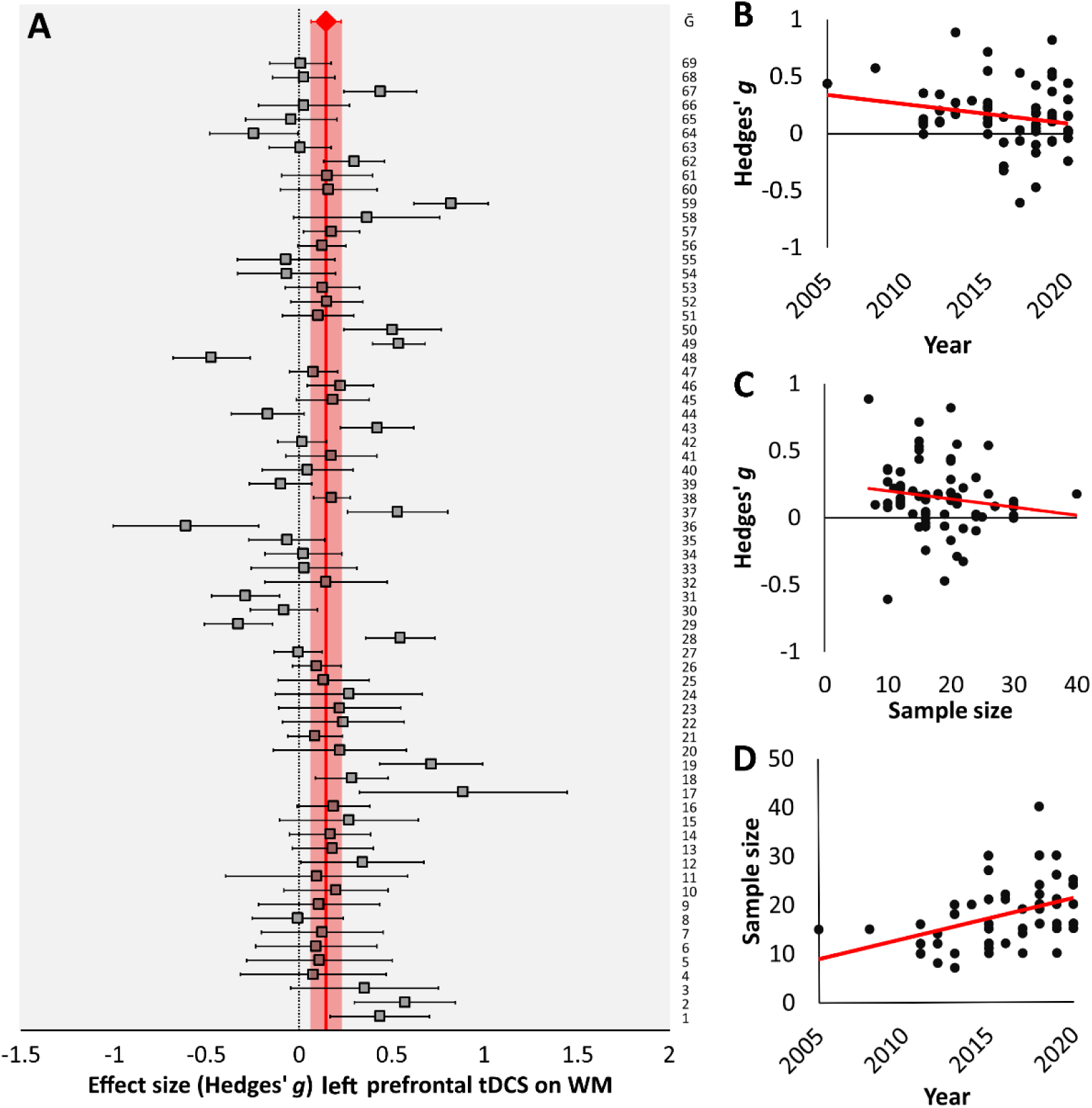
A) Forest plot showing effect sizes (Hedges’ g) and 95% confidence interval of left PFC anodal tDCS on working memory (n = 69). Positive values represent tDCS-related improved WM, whereas negative values represent decreased WM performance. Number labels on the right represent experiment numbers corresponding to Table 1. B) Scatterplots representing the relationship between publication year and effect size; C) between publication year and sample size; and D) between sample size and effect size.

The relationship between sample size, publication year and effect size were investigated to assess risk of publication bias. Publication year and effect size were not significantly correlated (ρ = −0.218, p = 0.072; Figure 2B). However, a slight negative trend was observed, which may be explained by the observation of a significant positive correlation between publication year and sample size (ρ = 0.482, p < 0.001; Figure 2C), as well as a significant negative correlation between sample size and effect size (ρ = −0.280, p = 0.020; Figure 2D). This suggests that more recent studies recruited more participants and studies with larger sample sizes tend to find smaller effect sizes. However, a funnel shape distribution and absence of asymmetry around the mean effect size suggest there is no clear evidence for publication bias (Figure 2D).

### 3.2. Classical meta-analysis for right prefrontal tDCS

Similar to the previous analysis, we performed a meta-analysis on the effect of right prefrontal tDCS on WM performance to get an indication of general tDCS efficacy irrespective of montage and intensity. For right PFC tDCS studies on WM performance 18 effect sizes were collected. Overall cumulative effect size did not reach significance (Ḡ = 0.115, 95% CI = −0.056 – 0.286, Z = 1.32, p = 0.187; Figure 3A). Total heterogeneity of effect sizes was not significant (Q_T_ = 6.95, p = 0.984) and Kolmogorov–Smirnov test indicated no deviation from normality (D = 0.204, p = 0.388). That is, without taking montage and intensity into account, right prefrontal tDCS is not effective in changing WM performance.

**Figure 3.**
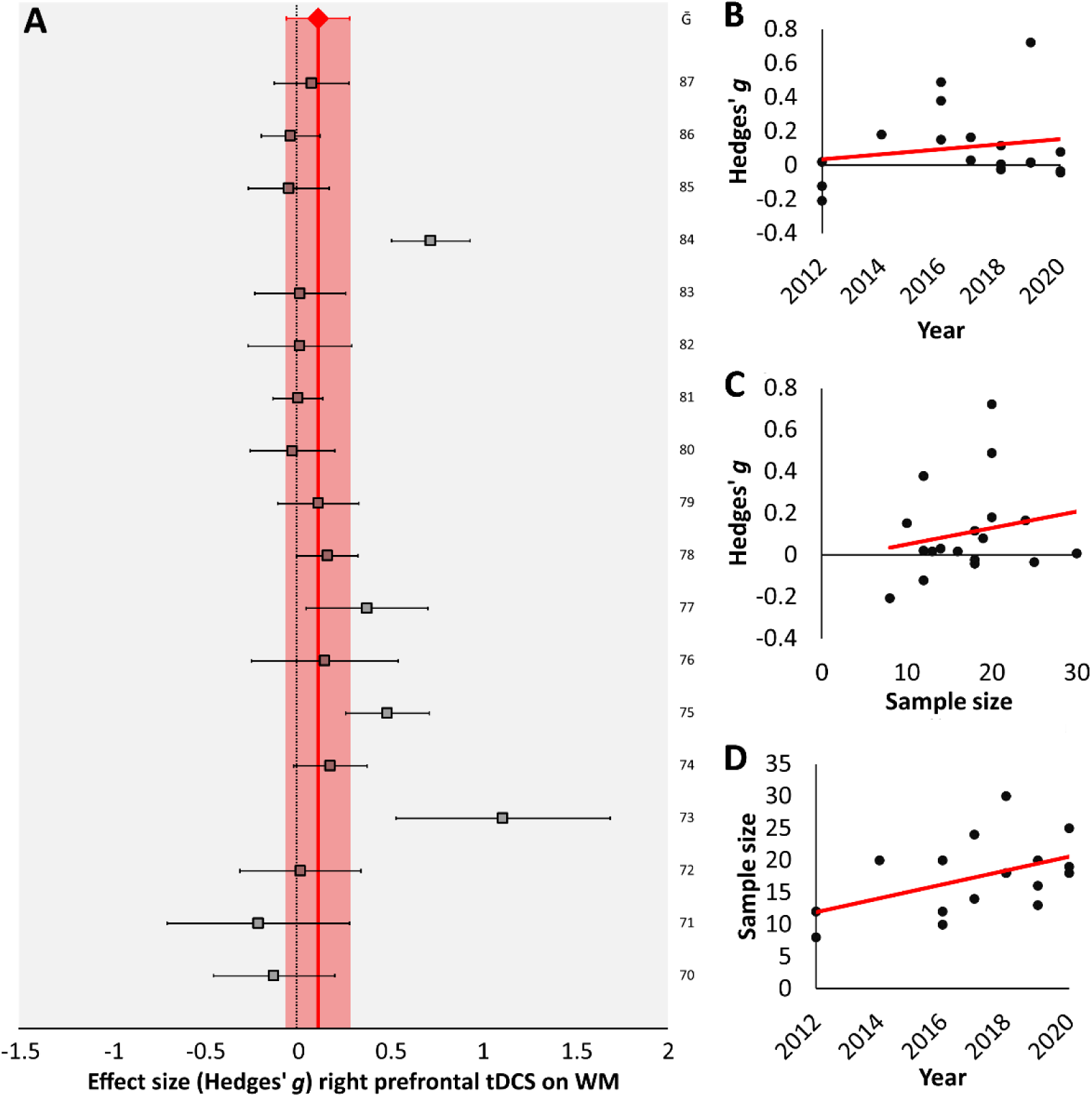
Forest plot showing effect sizes (Hedges’ g) and 95% confidence interval of right PFC anodal tDCS on working memory (n = 18). Positive values represent tDCS-related improved WM, whereas negative values represent decreased WM performance. Number labels on the right represent experiment numbers corresponding to Table 2. B) Scatterplots representing the relationship between publication year and effect size; C) between publication year and sample size; and D) between sample size and effect size.

**Table 2.**
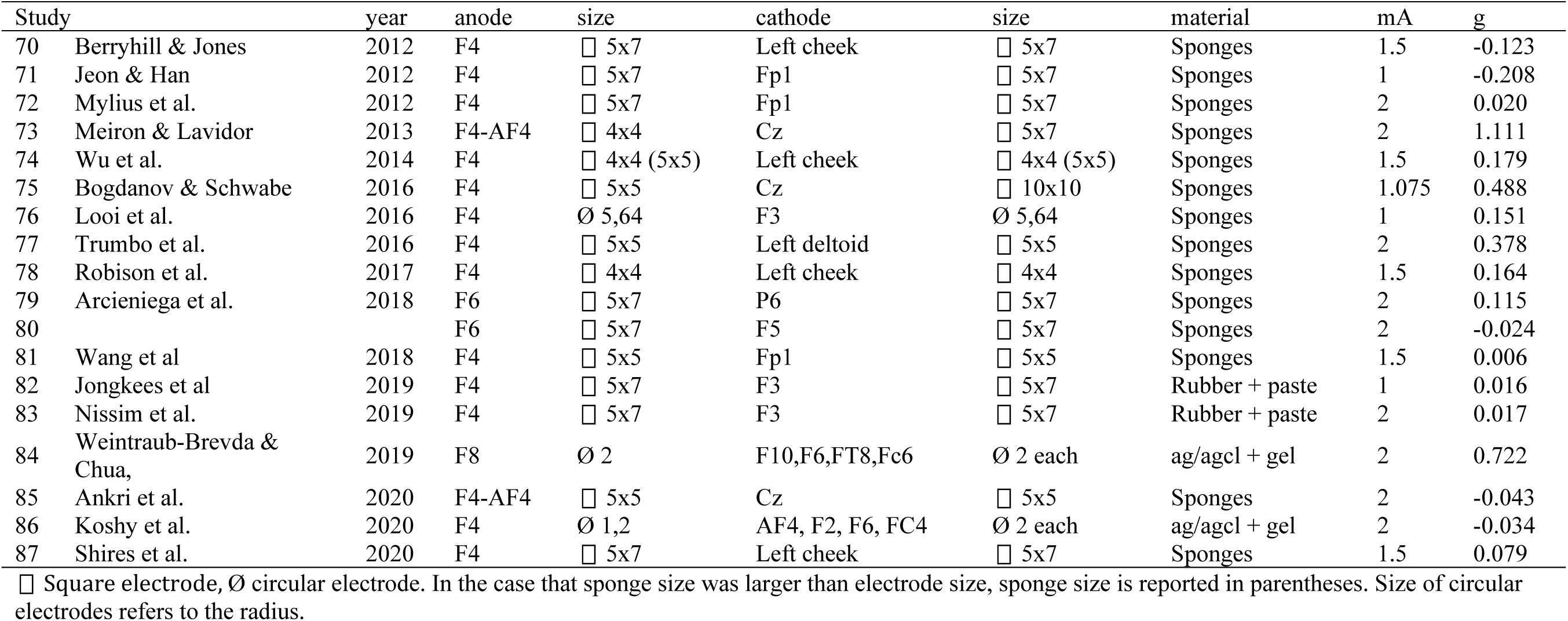
Study overview right PFC anodal tDCS

**Table 3.** Classical meta-analysis results

| | Montage | N | $\bar{G}$ | 95% CI | Z | p-value | Fail-safe number | Qt | p-value |
| --- | --- | --- | --- | --- | --- | --- | --- | --- | --- |
| Left PFC | Overall | 69 | 0.147 | 0.067 – 0.228 | 3.595 | < 0.001 | 306.5 | 40.48 | 0.997 |
|  | PFC-SOR | 28 | 0.141 | 0.010 – 0.273 | 2.101 | 0.036 | 30.4 | 13.23 | 0.988 |
|  | PFC-cheek | 4 | 0.208 | -0.454 – 0.869 | 0.615 | 0.539 | 0 | 0.09 | 0.993 |
|  | PFC-shoulder | 12 | 0.084 | -0.127 – 0.294 | 0.780 | 0.435 | 0 | 7.58 | 0.670 |
|  | PFC HD | 11 | 0.195 | -0.028 – 0.418 | 1.717 | 0.086 | 0 | 6.35 | 0.785 |
|  | PFC bifrontal | 6 | 0.018 | -0.355 – 0.392 | 0.095 | 0.924 | 0 | 3.66 | 0.599 |
|  | PFC other | 8 | 0.241 | -0.001 – 0.483 | 1.958 | 0.050 | 0 | 7.21 | 0.514 |
| Right PFC | Overall | 18 | 0.115 | -0.056 – 0.286 | 1.32 | 0.187 | 0 | 6.95 | 0.984 |

No significant relationship between publication year and effect size was observed (ρ = - 0.082, p = 0.745; Figure 3B). More recent publications were associated with larger sample sizes (ρ = 0.506, p = 0.032; Figure 3C), but no significant correlation between sample size and effect size was observed (ρ = 0.243, p = 0.331; Figure 3D). Furthermore, equal distribution around the mean suggests no publication bias (Figure 3D).

### 3.3. Left prefrontal relationship between electric field distribution and behavioral performance

Next, we performed our novel meta-analytic method on the correlation between electric field strength and behavioral performance. Initially, electric field distributions were simulated for all left prefrontal tDCS (n = 69) studies. The electric field averaged over all data suggested that a large portion of the PFC was targeted across studies and no clear preference for a specific region was observed (Figure 4A). The average Foc_50_ value of the 69 montages was 83.14 ± 5.56 cm^3^ and focality varied between 6.15 cm^3^ and 178.91 cm^3^, which corresponds to between 0.46% and 13.43% of total gray matter volume. This suggests that the extend of electric fields varied considerably across montages (Figure 4B-F, Supplementary Table 3). A significant difference between montage categories was observed (F(4,56) = 238.83, p < .001). Post-hoc analysis showed significant differences between all montages (p < .005), except between PFC-cheek and PFC bifrontal (p = .366). Unsurprisingly, average Foc_50_ was lowest for PFC HD (23.58 ± 3.83 cm^3^) and highest for PFC-shoulder (159.09 ± 4.46 cm^3^; Supplementary Table 2). E_MAX_ average was 0.307 ± 0.014 V/m and ranged between 0.113 V/m and 0.747 V/m and differed significantly between montage categories (F(4,56) = 3.23, p = 0.019). The only significant difference was between PFC-shoulder and PFC HD (p = 0.032).

**Figure 4.**
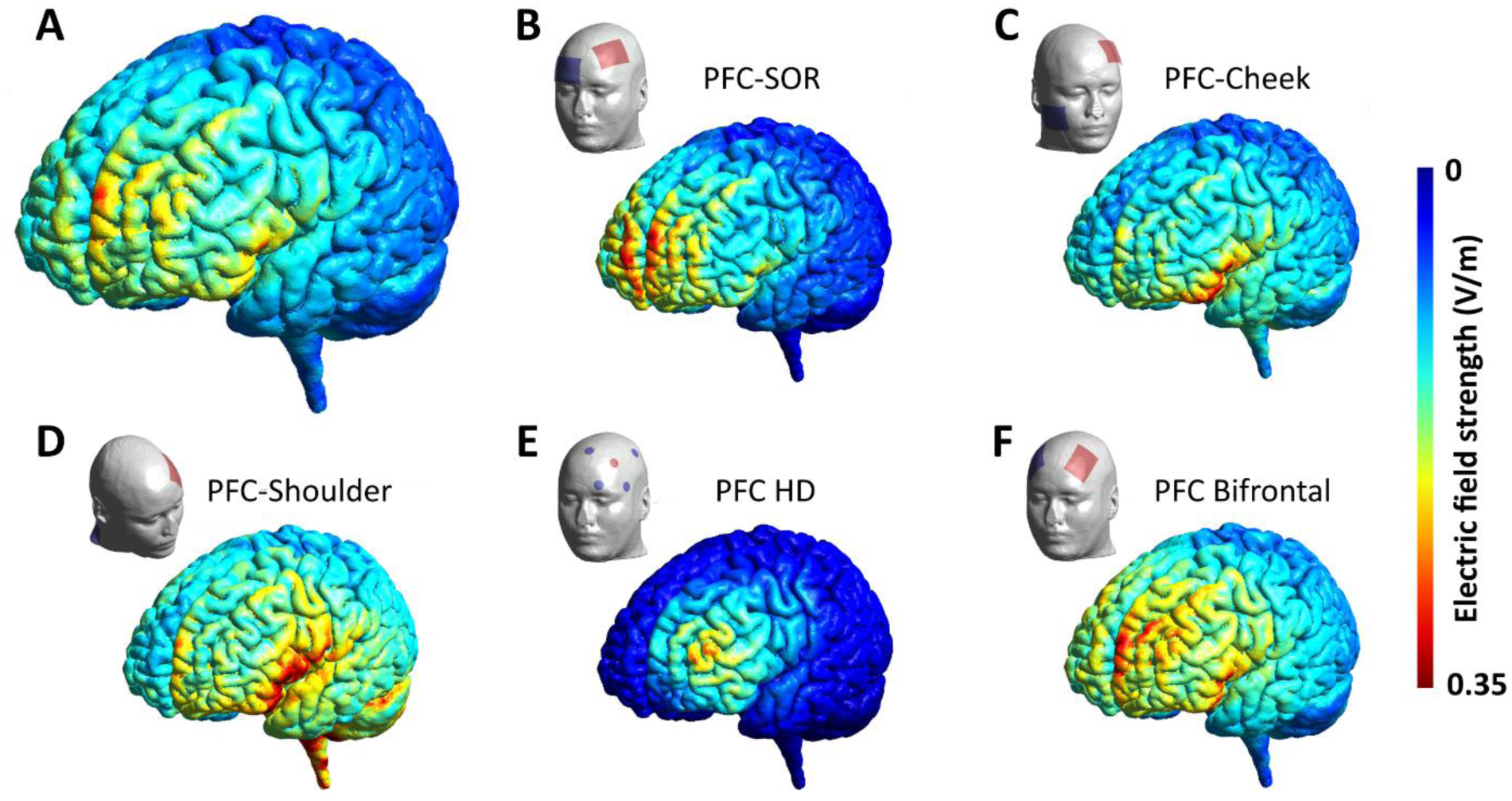
Averaged electric field distributions of left PFC tDCS. A) electric field strength averaged over all studies (n = 69). B) electric field strength averaged for PFC-SOR montages (n = 28); C) PFC-Cheek montages (n = 4); PFC-Shoulder montages (n = 12); PFC HD montages (n = 11); PFC bifrontal montages (n = 6).

Next, the relationship between electric field strength and behavioral effect size was investigated at each brain location. This relationship is represented by the PEC value, which is the correlation between electric field strength and Hedges’ g. We found a significant volume at the border of the lower left DLPFC and left IFC between Brodmann area 45 and 47 (Figure 5A). PEC_MAX_ was .279 (p = 0.010) at MNI coordinates [-51, 39, 4]. The PEC_MAX_ corresponds to 7.8% explained variance, which amounts to a medium-sized effect (Cohen, 1977). A total volume of 1.16 cm^3^ had a PEC value that reached significance (PEC > 0.199, p < 0.05; Figure 5B). Averaged PEC value within the significant volume was 0.241 (p = 0.023; Figure 6A).

**Figure 6.**
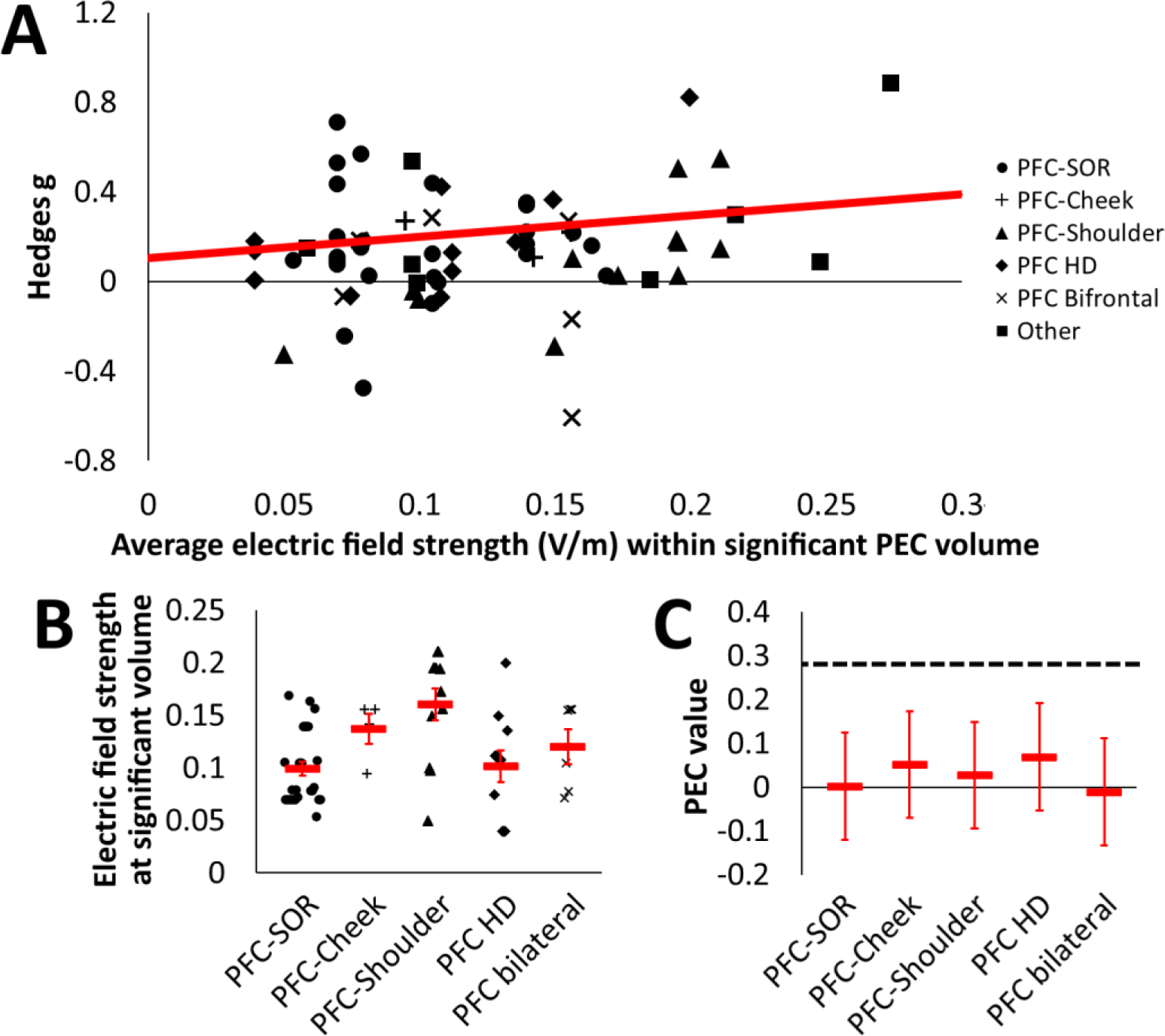
A) Scatterplot of the correlation between the electric field strength for each study averaged over the area of significant PEC values and effect size values. Various symbols are related to different montages. B) Individual and mean electric field strengths for each montage category averaged of the area of significant PEC values. C) PEC values at the E_MAX_ location of each montage category compared to PEC_MAX_ (dashed line).

We explored how well each of the five montage categories targets the region of significant PEC values. Electric field strength of each tDCS montage, averaged for all tetrahedra within the volume of significant PEC, differed considerably compared to their E_MAX_ (Figure 6B). This suggest that the PEC area was not primarily targeted by these montages. Furthermore, at the location that each montage targets the relationship to WM improvement is non-significant. Specifically, the PEC values corresponding to E_MAX_ location of each montage were not significantly larger than zero (Figure 6C): PFC-SOR [1, 62, 11], PEC = 0.002 (p = 0.990), PFC-Cheek [-32, 29, −15], PEC = 0.052 (p = 0.336), PFC-Shoulder [-30, −1, −24], PEC = 0.027 (p = 0.159), PFC HD [-39, 47, 23], PEC = 0.069 (p = 0.159), PFC Bifrontal [-23, 50, 15], PEC = - 0.011 (p = 0.159).

### 3.4. Right prefrontal relationship between electric field distribution and behavioral performance

The average Foc_50_ values of the 18 montages was 91.17 ± 10.52 cm^3^ and focality varied between 5.37 cm^3^ and 168.95 cm^3^, which corresponds to between 0.40% and 12.69% of the total gray matter volume. E_MAX_ average was 0.337 ± 0.027 V/m and ranged between 0.104 V/m and 0.509 V/m (Supplementary Table 3).

In agreement with the left PFC results, largest PEC values for tDCS over the right PFC were found in the lower part of the DLPFC (Figure 7A). Maximum PEC was 0.503 (p = 0.017), which corresponded to MNI coordinates [52, 36, 7]. A total volume of 1.29 cm^3^ had a PEC value that reached significance (PEC > 0.401, p < 0.05; Figure 7B).

### 3.5. Optimized tDCS montage

In order to induce electric fields with maximum strength in the significant PEC region, we propose a montage targeting the lower DLPFC/upper IFC. For this we used the electrode location optimization routine provided by SimNIBS. The routine resulted in a four-electrode montage with two circular (3.14 cm^2^) anodes over F7 and AF7, with an intensity of 1 mA each. Furthermore, two circular cathodes at an intensity of −1 mA over Fc1 and Fcz were suggested (Figure 8A). This montage yields an electric field over the significant PEC volume with E_MAX_ = 0.41 V/m and Foc_50_ = 56.39 cm^3^ (Figure 8B and C).

**Figure 8.**
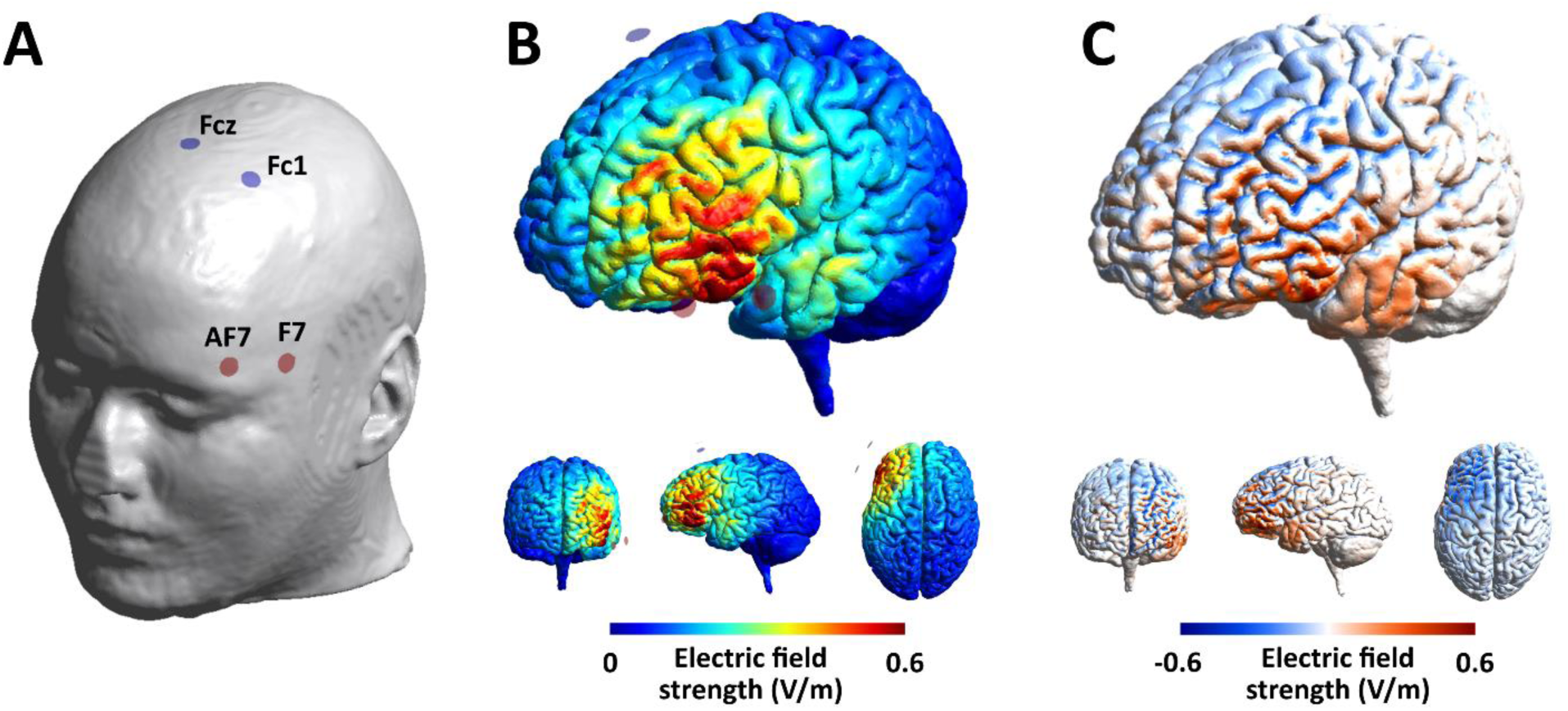
A) Montage resulting from the SimNIBS optimization routine to target an area with a radius of 10 mm around PEC_MAX_ in left PFC. Stimulation intensity for anodal electrodes at AF7 and F7 was 1 mA and for cathodal electrodes at Fc1 and Fcz was −1 mA. B) Induced electric field by the optimized montage. C) Inward (red) and outward (blue) currents induced by the optimized stimulation montage.

## Discussion

Here, we presented a novel meta-analytic approach, using computational electric field simulations to study the relationship between tDCS parameters and WM performance. Large variety in tDCS set ups yields large inter-experimental variability in electric field distribution and strength (Opitz et al., 2018; Polania et al., 2018). As a result, classical meta-analyses are not well-suited to capture differences between studies using different tDCS montages. Instead, our method exploited the observed variability in tDCS electric fields to predict brain regions where tDCS is related to changes in WM performance. Analysis related to 69 effect sizes of anodal tDCS over left PFC showed that tDCS electric fields in the lower left DLPFC/upper left IFC (Brodmann area 45 and 47) are most strongly related to increased general WM performance. Differences in electric field strength in this region accounts for 7.8% of variance for WM improvements, which amounts to a medium-sized effect (Cohen, 1977). Interestingly, a similar region for tDCS-induced WM performance improvements was observed for right PFC tDCS studies (n = 18). Electric fields of five tDCS montages categories commonly used in WM-related research did not reach maximum strength in the lower DLPFC/upper IFC. Therefore, we presented a tDCS montage to better target this region. Hence, future studies could investigate the potential performance benefits on WM performance of this tDCS montage.

### 4.1. Electric field modeling on meta-analytic datasets

In the present study we introduced a novel way of systematically investigating tDCS parameters across multiple studies. Recently, studies used individualized electric field simulations and compared these to changes in behavior (Albizu et al., 2020; Caulfield et al., 2020; Evans et al., 2020; Kim et al., 2014). Thereby, these studies were able to control for inter-individual electric field variability when interpreting tDCS-related effects. Analogously, here inter-experiment electric field variability was exploited to explain differences between studies in tDCS-induced WM performance change. Effects of non-invasive brain stimulation are prone to numerous sources of intra-individual, inter-individual, and inter-experimental variability (Lopez-Alonso et al., 2014; Polania et al., 2018; Ridding & Ziemann, 2010). Although variability should be reduced, it cannot be completely eliminated. With our new approach, we were able to overcome shortcomings of classical meta-analyses, which are unable to tackle variability across studies with respect to differences to tDCS stimulation parameters. As such, we calculated a brain region specific estimate of tDCS-related WM improvements. Indeed, whereas previous meta-analyses either found small or null effects of tDCS on WM (Brunoni & Vanderhasselt, 2014; Hill et al., 2016; Horvath et al., 2015; Manusco et al., 2016), our method revealed that a medium effect (7.8% explained variance), specifically for the lower DLPFC/upper IFC region. In sum, converging evidence of 87 effect sizes, reported in 68 peer-reviewed articles, suggested that anodal tDCS can improve WM performance and that targeting the lower DLPFC may be the best way to achieve this.

### 4.2. Differences in PFC tDCS target region

Investigation of electric field distributions related to different tDCS montages showed considerable variation in targeted volume. Orbital-referenced and bifrontal montages primarily target anterior portions of the PFC, corresponding to Brodmann area 10. In contrast, cheek- and shoulder-referenced montages target more posterior and inferior regions, such as triangular and opercular part of the IFC, corresponding to Brodmann areas 44 and 45. High-definition montage electric field distribution is more restricted to location of the anodal montages. Since most studies used F3 of the 10-20 system for electrode positioning, the induced electric field is largest in the superior part of the DLPFC (Brodmann area 46). The present meta-analytic results suggested that the inferior part of the DLPFC and superior orbital part of the IFC were most strongly related to increased WM performance. Electric fields induced by five common montages were not maximal at this region, providing a potential explanation for overall small effect sizes on WM performance.

Indeed, the majority of tDCS studies intended to target the upper DLPFC by placing the anodal electrode over F3. However, a few studies attempted to target lower DLPFC/IFC by placing the target electrode at F5/F6 or F7/F8 (Arciniega et al., 2019; Di Rosa et al., 2019; Jones et al., 2015; Lukasik et al., 2018; Weintraub-Brevda & Chua, 2019). For example, Weintraub-Brevda and Chua (2019) found that anodal tDCS over left and right IFC (referred to as ventrolateral PFC) increased both emotional and neutral WM. Similarly, Jones et al. (2015) found WM improvements of lower PFC anodal tDCS, but these results were limited to participants with high baseline WM capacity. Furthermore, tDCS-effects were amplified with the presentation of extrinsic motivation by offering a reward. These results were partially confirmed by Di Rosa et al. (2019), who showed lower PFC anodal tDCS-related improved reaction time during and after a reward-driven WM task. However, in contrast to Jones et al., Di Rosa and colleagues found that the effect was stronger for participants with lower baseline WM capacity. Furthermore, in agreement with our findings, the beneficial tDCS-induced effects on WM were associated with increased hemodynamic activity in the IFC. Contrary to these findings, Lukasik et al. (2018) reported no tDCS-related effects on WM when the anodal electrode was placed over IFC. Yet, whereas the montages used by Jones et al. (2015), Di Rosa et al. (2019), and Weintraub-Brevda & Chua (2019) electric field distribution significantly overlapped with the IFC (Supplementary Figure 2, study nr. 22, 51, and 59), Lukasik et al. (2018) targeted more anterior parts of the PFC (Supplementary Figure 2, study nr. 47). This further demonstrates the benefits of electric field modeling when determining affected cortical volumes.

Although the mid-to-upper DLPFC has received much attention and was targeted by most tDCS studies on WM performance, considerable evidence supports a crucial role for the lower DLPFC/upper IFC (Grossberg, 2018; Petrides et al., 2002). Various non-mutually exclusive hypotheses have been proposed that point towards different functional properties of upper and lower PFC, including manipulation and maintenance of information (Petrides et al., 2002; Stern et al., 2000; Veltman et al., 2003), distinct processing of neutral and emotional stimuli (Dolcos et al., 2013), and a spatial versus verbal distinction (Grossberg, 2018; Owen et al., 1999). Furthermore, within PFC areas, different neuronal activity patterns have been observed. Whereas some neurons display elevated activity during a delay period, others show an ‘activity-silent’ pattern where neurons maintain information by a change in synaptic weights (Spaak et al., 2017; Stokes, 2015). Future research may investigate whether tDCS-induced cortical activation might preferentially target specific functional processes and neuronal encoding patterns and how this relates to different PFC subdivisions.

### 4.3. Potential biases in WM-related tDCS studies

Our risk of bias assessment suggested no strong biases within studies. Blinding of experimenters was not guaranteed in one third of studies, which poses a small potential of performance bias. However, report of successful blinding of participants, as well as their sensations during active tDCS and sham is frequently lacking and should be adopted in the future. Overall, we found no strong indication for publication bias (Sterne et al., 2011). Yet, it should be noted that more recent studies tended to recruit a larger sample size, which is associated with smaller effect sizes. Indeed, larger sample sizes are associated with smaller variance and are thus less likely to include extremely large (or small) effects. In accordance, although not statistically significant, we found a downward trend in effect sizes correlated to publication year. This observation may be related to a recent increase in openness to publish null findings (Schäfer & Schwarz, 2019).

### 4.4. Limitations and perspectives

One limitation of our study relates to the variability in tasks and outcome measurements between studies. We opted to include several WM tasks (N-back, Sternberg task, digit span, etc.), as well as outcome measurements (accuracy, RT, achieved N, etc.), in order to present results of tDCS effects that can be extrapolated over all WM-related tasks. Investigating specific aspects of WM could have reduced inter-experimental variability, at the cost of generalizability. Furthermore, selecting specific parameters of a task would have reduced sample size, which would reduce statistical power.

Furthermore, it should be noted that although the present study accounts for inter-experimental variability, inter-individual variability is an additional determinant for tDCS efficacy (Evans et al., 2020). Here, we used a typically healthy male brain for electric field modeling. However, head shape, brain size, skull-to-cortex distance, gyrification and conductivity differs between individuals (Antonenko et al., 2021; Opitz et al., 2015). Consequently, the present results are generalizable to a group-level, however precise location related to optimal tDCS-effects on WM can differ per individual. Therefore, adjusting tDCS montage based on individual imaging data is desirable (Evans et al., 2020). This issue is further amplified when considering patient population with brain atrophy or brain damage (Lu et al., 2019). Whether present results translate to abnormal brain physiology needs to be established in future studies.

Although electric field distribution varied considerably between studies, systematic experimental investigation of montage configuration, as well as target location may be a subject for future studies. Comparison of tDCS that primarily targets IFC, lower DLPFC, upper DLPFC and anterior PFC offers the opportunity to experimentally verify current meta-analytic findings. Furthermore, the efficacy of our tDCS montage suggestion, which followed from the optimization routine, should be empirically substantiated.

Besides tDCS, applying oscillatory currents using transcranial alternating current stimulation (tACS) has been proposed as a method of changing neural activity and behavioral performance (Johnson et al., 2020; Neuling et al., 2013; Reato et al., 2013; Schutter & Wischnewski, 2016; Wischnewski et al., 2019a; 2019b). Recent studies have suggested that tACS can improve WM performance when targeting theta oscillations (Abellaneda-Perez et al., 2020, Jausovec & Jausovec, 2014; Jausovec et al., 2014; Röhner et al., 2018), gamma oscillations (Hoy et al., 2015), or a combination of both (Alekseichuk et al., 2016). Interestingly, Abellaneda-Perez and colleagues (2020) suggested differing neural network activation related to tDCS- and tACS-related effects on WM performance. Indeed, neural mechanisms of tACS may differ from tDCS, where neurons are entrained without an increase in spiking rate (Johnson et al., 2020). As such, the differing neurophysiological properties that tDCS and tACS may target mean that the translation of our findings to tACS applications needs to be established.

WM deficits have been observed in a variety of disorders, including attention-deficit hyperactivity disorder (ADHD) (Salehinejad et al., 2020; Valera et al., 2005), post-traumatic stress disorder (PTSD) (Flanagan et al., 2018), obsessive-compulsive disorder (OCD) (Nakao et al., 2009), and major depressive disorder (MDD) (Joormann & Gotlib, 2008; Marquand et al., 2008). Findings of our meta-analysis may provide insights for clinical studies that employ tDCS to treat individuals suffering from these and other disorders. Indeed, tDCS has been shown to improve WM performance in children with ADHD, however, the effect was small and large variability was observed (Salehinejad et al., 2019). Similarly, several studies provided evidence for tDCS-related WM benefits in patients with MDD (Moreno et al., 2015; Oliveira et al., 2013), which has been associated with relief of depressive symptoms (Salehinejad et al., 2015). Therefore, attempting to maximize tDCS-related effects to WM by performing meta-analytic electric field modeling studies, as we propose here, may be beneficial to the therapeutic use of tDCS in the treatment of psychiatric disease.

