## supplementary material for "Identifying regions in prefrontal cortex related to working memory improvement: a novel meta-analytic method using electric field modeling"

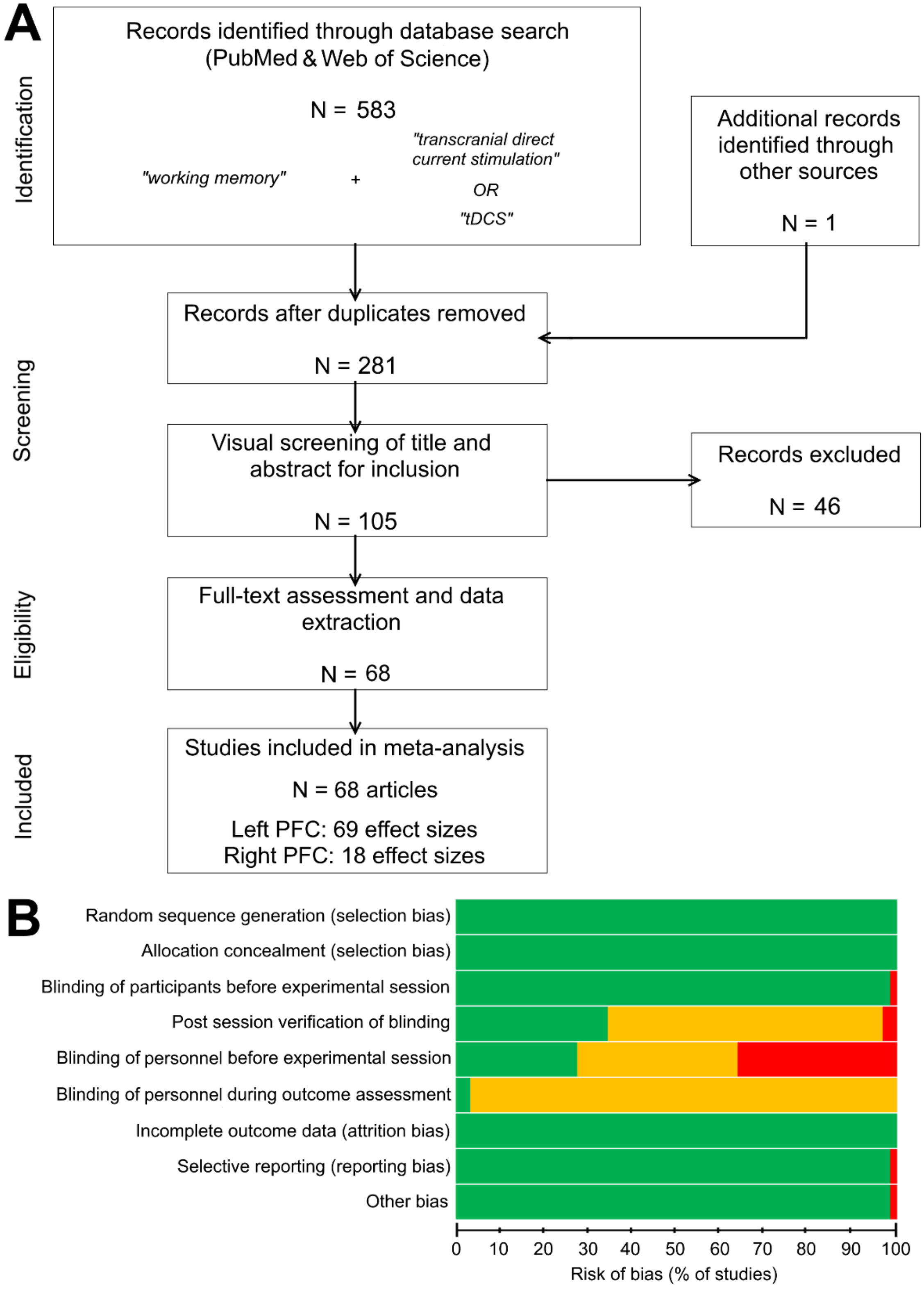

**Supplementary Figure 1**. A) Preferred Reporting Items for Systematic Reviews and Meta-Analyses (PRISMA) flow diagram of selection of studies. B) Risk of bias assessment. Overall risk of bias was low. Specifically, assessment suggested low risk of selection bias, attrition bias, reporting bias, performance bias related to blinding of participants, and other uncategorized biases. However, post experimental verification of participant blinding and blinding of personnel during outcome assessment was not reported in most studies. Risk of performance bias related to blinding of personnel during the experiment was observed, with a third of studies using a single-blind design and another third of studies not reporting blinding of personnel.

**Supplementary Table 1** Demographics, task overview, outcome measure, control, stimulation duration and timing of left prefrontal tDCS studies

| Study | | year | N (f/m) | Age ± SEM | Task | Outcome | Control | Duration | Online/ |
|---|---|---|---|---|---|---|---|---|---|
| 1 | Fregni et al. | 2005 | 15 (11/4) | 20.2 (19-22) | 3-back | Accuracy & RT | Sham | 10 | On |
| 2 | Ohn et al. | 2008 | 15 (10/5) | 26.5 ± 0.9 | 3-back | Accuracy & RT | Sham & BC | 30 | On & Off |
| 3 | Andrews et al. | 2011 | 10 (6/4) | 28.1 ± 2.8 | Digit span | Span forward & | Sham & BC | 10 | Off |
| 4 | Keeser et al. | 2011 | 10 (5/5) | 28.9 ± 0.8 | 2-back | Accuracy & RT | Sham[1] | 20 | Off |
| 5 | Mulquiney et al. | 2011 | 10 (6/4) | 29.4 ± 1.8 | 2-back | Accuracy & RT | Sham & BC | 10 | Off |
| 6 | Teo et al. | 2011 | 12 (7/5) | 27.2 ± 2.7 | Sternberg & 3-back | Accuracy & RT | Sham | 20 | On & Off |
| 7 | | | | | | | | | |
| 8 | Zaehle et al.[2] | 2011 | 16 (10/6) | 25 ± 0.5 | 2-back | Sensitivity & RT | Sham & CG | 15 | Off |
| 9 | Berryhill & Jones | 2012 | 24 (?) | 63.7 (56-80) | 2-back | Accuracy | Sham | 10 | Off |
| 10 | Gladwin et al. | 2012 | 14 (8/6) | 22 ± 0.8 | Sternberg task | Accuracy & RT | Sham | 10 | On |
| 11 | Jeon & Han | 2012 | Real: 8 (3/5) | 39.5 ± 5.3 | Digit span | Span forward & | Sham & BC | 20 | Off |
| 12 | Mylius et al. | 2012 | 12 (6/6) | 25.1 ± 3.4 | 2-back | Accuracy & RT | Sham | 20 | On |
| 13 | Hoy et al. | 2013 | 18 (11/7) | 24.7 ± 1.6 | 2-back & 3-back | Accuracy & RT | Sham | 20 | Off |
| 14 | | | | | | | | | |
| 15 | Lally et al. | 2013 | Real: 10 (?) | 23.1 ± 0.9[3] | 3-back | Sensitivity & RT | Sham & BC | 10 | On & Off |
| 16 | Martin et al. | 2013 | Real: 14 (9/12) | 23.2 ± 1.5 | Individually adjusted | Sensitivity | Sham & BC | 30 | On |
| 17 | Meiron & Lavidor | 2013 | Real: 14 (7/7) | 24.9 ± 0.8 | 2-back | Accuracy | Sham | 15 | On |
| 18 | Richmond et al. | 2014 | Real: 20 (13/7) | 20.7 ± ? | Verbal & spatial WM | Span | Sham | 15 | On |
| 19 | Carvalho et al. | 2015 | 15 (12/3) | 20.2 ± 0.7 | 3-back | Accuracy | Sham | 20 | On |
| 20 | Gill et al. | 2015 | 12 (5/7) | 19.8 ± 0.4 | 3-back & adjusting | Accuracy | Sham & BC | 20 | On & Off |
| 21 | Hussey et al. | 2015 | Real: 27 (12/15) | 19.5 (18-23) | 2-back & 4-back | Discriminability | Sham | 30 | On & Off |
| 22 | Jones et al. Exp 1 | 2015 | 24 (12/12) | 23.8 ± 0.7 | Change detection task | Accuracy | Sham | 10 | Off |
| 23 | Jones et al. Exp 2 | 2015 | 20 (12/8) | 22.0 ± 0.7 | Change detection task | Accuracy | Sham | 10 | Off |
| 24 | Moreno et al. | 2015 | Real: 10 (5/5) | 26.3 ± 2.5 | 2-back & internal shift | Accuracy, RT & | Sham | 30 | Off |
| 25 | Nikolin et al. | 2015 | 16 (8/8) | 21.8 ± 0.6 | 3-back | Accuracy & RT | Sham | 20 | Off |
| 26 | Nilsson et al. | 2015 | 30 (14/16) | 69 ± 1.3 | 3-back | Accuracy & RT | Sham & BC | 25 | On & Off |
| 27 | | | | | | | | | |
| 28 | Pope et al. | 2015 | Real: 21 (13/8) | 22.0 ± 1.1 | PASAT & PASST | Accuracy & RT | Sham | 20 | Off |
| 29 | Faehling & Plewnia | 2016 | Real: 22 (22/0) | 23.9 ± 0.3[4] | Delayed WM task | Accuracy & RT | Sham | 28 | On |
| 30 | | | Real: 22 (22/0) | | | | | | |
| 31 | | | Real: 21 (21/0) | | | | | | |
| 32 | Trumbo et al. | 2016 | Real: 12 (6/6) | 20.5 ± 1.0 | Spatial & verbal 3-back | Accuracy, RT & | Sham & BC | 30 | On & Off |
| 33 | Cespon et al. | 2017 | Young: 14 (6/8) | 24.8 ± 1.0 | 2-back & 3-back | Sensitivity & RT | Sham & BC | 13 | Off |
| 34 | Hill et al. | 2017 | 20 (12/8) | 29.1 ± 2.8 | 2-back | Sensitivity & RT | Sham & BC | 20 | Off |
| 35 | | | | | | | | | |
| 36 | Nikolin et al. | 2017 | Real: 10 (4/6) | 22.3 ± 1.2 | 3-back | Accuracy & RT | Sham & BC | 15 | On & Off |
| 37 | Talsma et al. | 2017 | Real: 15 (11/4) | 21.9 ± 0.7 | 3-back and 4-back | Accuracy | Sham & BC | 20 | On & Off |
| 38 | Deldar et al. | 2018 | 40 (23/17) | 25.8 ± 0.7 | 2-back | Accuracy & RT | Sham & BC | 22 | On |
| 39 | Dumont et al. | 2018 | 24 (14/10) | 21.1 ± 0.4 | NIH-examiner WM | WM score | Sham | 20 | Off |
| 40 | Hill et al. | 2018 | 16 (10/6) | 32.8 ± 2.7 | 2-back | Sensitivity & RT | Sham & BC | 15 | Off |
| 41 | | | | | | | | | |
| 42 | Lukasik et al. | 2018 | 33 (23/10) | 22.6 ± 0.4 | 3-back | Sensitivity & RT | Sham & BC | 10 | On & Off |
| 43 | Naka et al. | 2018 | 20 (10/10) | 22.7 ± 0.9 | Visual & Auditory 3- | Accuracy & RT | Sham & BC | 16 | On & Off |
| 44 | Nikolin et al. | 2018 | Real: 20 (?) | 22.9 ± 0.4[4] | 3-back | Sensitivity & RT | Sham & BC | 15 | On & Off |
| 45 | | | | | | | | | |

| | | | | | | | | | |
|---|---|---|---|---|---|---|---|---|---|
| 46 | Rabipour et al. | 2018 | Real: 45 (27/18) | 20.5 ± 0.4[4] | 3-back | Accuracy | Sham | 20 | On |
| 47 | Rohner et al. | 2018 | 30 (15/15) | 26.2 ± 0.5 | 2-back | Accuracy & RT | Sham & BC | 15 | On & Off |
| 48 | Talsma et al. | 2018 | 20 (0/20) | 21.8 ± 0.6 | 3-back and 4-back | Accuracy & RT | Sham & BC | 20 | On & Off |
| 49 | Baumert et al. | 2019 | Real: 31 (?) | 23.0 ± 0.6[3] | 2-back & 3-back | Accuracy & RT | Sham | 20 | Off |
| 50 | Deldar et al. | 2019 | 15 (7/8) | 64 ± 1.1 | 2-back | Accuracy & RT | Sham & BC | 22 | On |
| 51 | Di Rosa et al. | 2019 | 21 (12/9) | 69.7 ± 1.1 | Custom spatial WM | Accuracy & RT | Sham & BC | 26 | On & Off |
| 52 | Friehs & Frings | 2019 | Real: 21 (13/8) | 24.2 ± 0.5 | 3-back | Accuracy & RT | Sham & BC | 19 | On & Off |
| 53 | Hill et al. | 2019 | 20 (11/9) | 24.1 ± 1.8 | 2-back | Sensitivity & RT | Sham & BC | 15 | Off |
| 54 | Jongkees et al | 2019 | Real: 48 (32/16) | 22.0 ± 0.7 | 2-back & 4-back | Sensitivity & RT | Sham & BC | 15 | Off |
| 55 | Ke et al. | 2019 | Real: 15 (?) | 20-25[3] | N-back, 3-back & 4- | Achieved N & | Sham & BC | 25 | On |
| 56 | Luque-Casado et al. | 2019 | 30 (7/23) | 21.6 ± 0.5 | Digit span | Backward span | Sham & BC | 15 | Off |
| 57 | Nikolin et al. | 2019 | Real: 26 (18/8) | 22.6 ± 0.9 | 2-back | Accuracy & RT | Sham | 12 | On & Off |
| 58 | Wang et al. | 2019 | Real: 10 (?) | 23.2 ± 0.4[3] | N-back | Achieved N | Sham & BC | 30 | On |
| 59 | Weintraub-Brevda | 2019 | Real: 20 (?) | 18-35[3] | Delayed WM task | Accuracy | Sham | 20 | On |
| 60 | Abellaneda-Perez et | 2020 | Real: 15 (7/8) | 24.3 ± 1.1 | 2-back & 3-back | Sensitivity & RT | Sham | 19 | On & Off |
| 61 | Byrne et al. | 2020 | Real: 16 (9/7) | 23.2 ± 0.9 | Digit span | Backward span | Sham & BC | 10 | On |
| 62 | Hussey et al. | 2020 | Real: 24 (12/12) | 23.1 ± ? | 2-back & 4-back | Discriminability | Sham & BC | 30 | On |
| 63 | Koshy et al. | 2020 | 25 (10/15) | 23.4 ± 0.7 | Visual WM task | Accuracy & RT | Sham | 20 | Off |
| 64 | Murphy et al. | 2020 | Real: 16 (11/5) | 30.4 ± 3.0 | Sternberg task | Accuracy & RT | Sham & BC | 22 | Off |
| 65 | Papazova et al. | 2020 | 1 mA: 16 (7/9) | 31.1 ± 0.9 | 2-back & 3-back | Sensitivity & RT | Sham | 21 | On |
| 66 | | | 2 mA: 16 (4/12) | 32.1 ± 2.0 | | | | | |
| 67 | Ramaraju et al. | 2020 | 20 (0/20) | 30 ± 1.8 | 2-back | Accuracy | Sham | 15 | Off |
| 68 | Splittgerber et al. | 2020 | 24 (13/11) | 24.8 ± 0.6 | 2-back | Accuracy & RT | Sham | 20 | On & Off |
| 69 | | 2020 | | | | | | | |

Abbreviations: BC = baseline control, CG = Control group, Off = offline, On = online, PASAT = paced auditory serial addition task, PASST = paced auditory serial subtraction

**Supplementary Table 2**. Demographics, task overview, outcome measure, control, stimulation duration and timing of right prefrontal tDCS studies

| Study | | year | N (f/m) | Age ± SEM | Task | Outcome | Control | Duration | Online/ |
|---|---|---|---|---|---|---|---|---|---|
| 70 | Berryhill & Jones | 2012 | 24 (?) | 63.7 (56-80) | 2-back | Accuracy | Sham | 10 | Off |
| 71 | Jeon & Han | 2012 | Real: 8 (4/4) | 35.1 ± 4.2 | Digit span | Span forward & | Sham & BC | 20 | Off |
| 72 | Mylius et al. | 2012 | 12 (10/2) | 23.5 ± 1.1 | 2-back | Accuracy & RT | Sham | 20 | On |
| 73 | Meiron & Lavidor | 2013 | Real: 16 (8/8) | 25.4 ± 1.4 | 2-back | Accuracy | Sham | 15 | On |
| 74 | Wu et al. | 2014 | 20 (8/12) | 26 (24-31) | Corsi block tapping | Span forward & | Sham | 15 | On |
| 75 | Bogdanov & | 2016 | Real: 20 (10/10) | 25.3 ± 0.3[1] | Corsi block tapping | Span backward | Sham & BC | 6-10 | On |
| 76 | Looi et al. | 2016 | Real: 10 (6/4) | 24.6 ± 1.2 | Corsi block tapping | Span forward & | Sham | 30 | On |
| 77 | Trumbo et al. | 2016 | Real: 12 (6/6) | 20.2 ± 0.9 | Spatial & verbal 3-back | Accuracy, RT & | Sham & BC | 30 | On & Off |
| 78 | Robison et al. | 2017 | 24 (?) | ? | Change detection task | WM capacity & | Sham | 20 | On & Off |
| 79 | Arcieniega et al. | 2018 | Young: 36 (22/14) | 21.2 ± 0.4 | Custom Visual WM | Accuracy | Sham | 20 | On |
| 80 | | | | | | | | | |
| 81 | Wang et al | 2018 | Real: 10 (?) | 23.2 ± 0.4[1] | N-back | Achieved N | Sham & BC | 30 | On |
| 82 | Jongkees et al | 2019 | Real: 40 (24/16) | 22.2 ± 0.8 | 2-back & 4-back | Sensitivity & RT | Sham & BC | 15 | Off |
| 83 | Nissim et al. | 2019 | 16 (6/10) | 71.75 ± 1.8 | 2-back | Accuracy & RT | Sham & BC | 12 | On & Off |
| 84 | Weintraub-Brevda | 2019 | Real: 20 (?) | 18-35[1] | Delayed WM task | Accuracy | Sham | 20 | On |
| 85 | Ankri et al. | 2020 | Real: 35 (27/8) | 23.9 ± 0.2[1] | 2-back | Accuracy | Sham | 20 | On |
| 86 | Koshy et al. | 2020 | 25 (10/15) | 23.4 ± 0.7 | Visual WM task | Accuracy & RT | Sham | 20 | Off |
| 87 | Shires et al. | 2020 | Real: 19 (?) | 22.6 ± 0.6[1] | Change detection task | Accuracy | Sham | 15 | On |

Abbreviations: BC = baseline control, Off = offline, On = online, RT = reaction time, WM = working memory

tDCS to left PFC

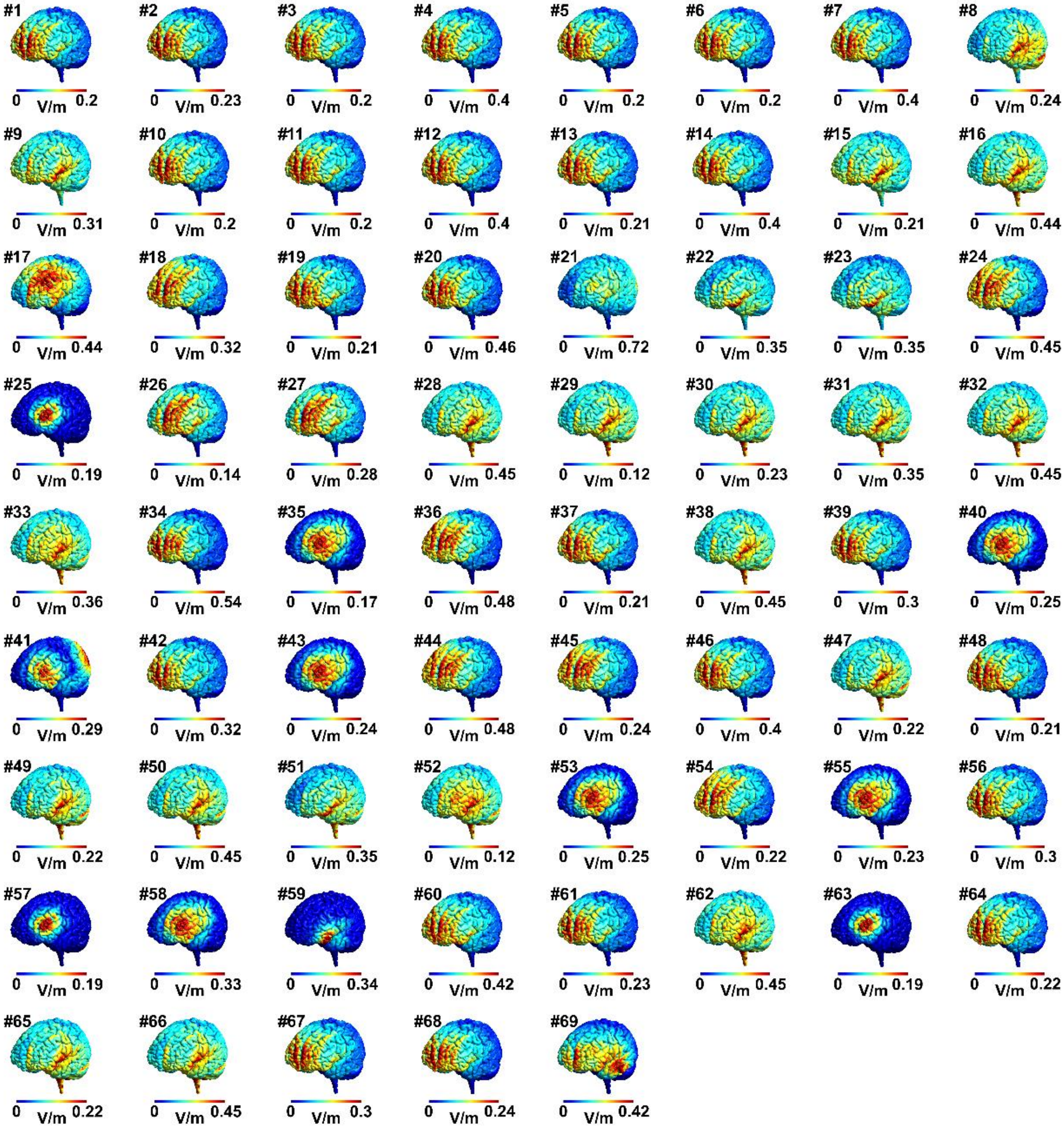


tDCS to right PFC

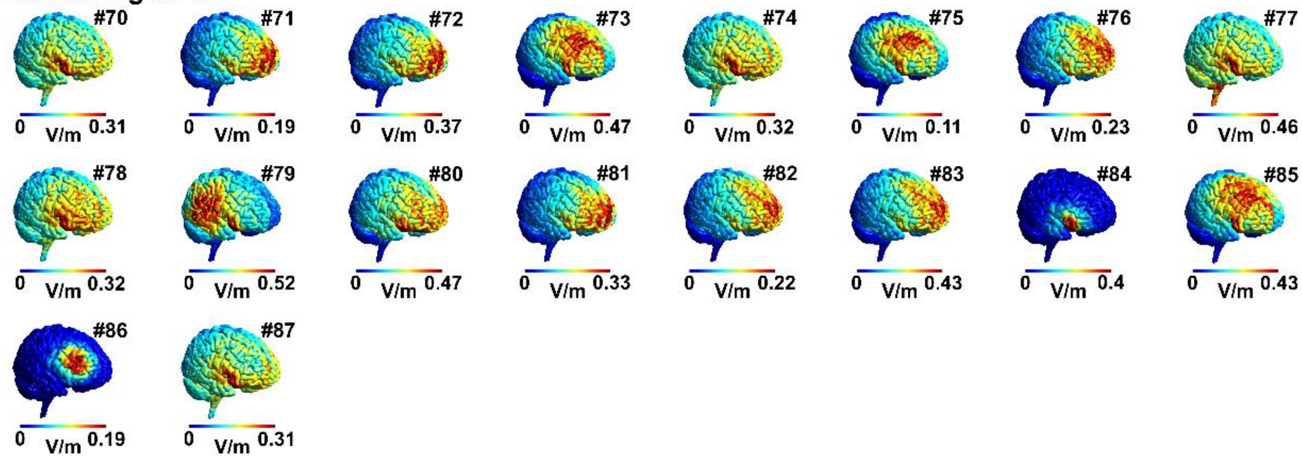

**Supplementary Figure 2**. Electric field distribution for all studies included in the present meta-analysis. Montage number correspond to study numbers of Table 1 and 2, as well as Supplementary Table 1 and 2.

**Supplementary table 3.** Electric field strength and focality.

| | | $E_{MAX}$ (V/m) | $E_{95}$ (V/m) | $E_{75}$ (V/m) | $E_{50}$ (V/m) | $Foc_{50}$ ($cm^3$) | $Foc_{50}$ ($\%_{TOTAL}$) | $Foc_{75}$ ($cm^3$) | $Foc_{75}$ ($\%_{TOTAL}$) |
|---|---|---|---|---|---|---|---|---|---|
| All left | M ± SEM | 0.307 ± 0.014 | 0.177 ± 0.009 | 0.103 ± 0.007 | 0.069 ± 0.006 | 83.14 ± 5.56 | 6.24 ± 0.42 | 10.08 ± 0.34 | 0.76 ± 0.03 |
| | Min-Max | 0.113-0.747 | 0.033-0.342 | 0.009-0.241 | 0.004-0.191 | 6.15-178.91 | 0.46-13.43 | 2.49-14.65 | 0.19-1.10 |
| PFC-SOR | M ± SEM | 0.292 ± 0.020 | 0.171 ± 0.011 | 0.090 ± 0.006 | 0.053 ± 0.003 | 66.28 ± 0.85 | 4.98 ± 0.06 | 10.65 ± 0.14 | 0.80 ± 0.011 |
| | Min-Max | 0.142-0.545 | 0.083-0.301 | 0.047-0.140 | 0.030-0.082 | 51.57-74.05 | 3.87-5.56 | 9.47-13.40 | 0.71-1.01 |
| PFC-cheek | M ± SEM | 0.298 ± 0.029 | 0.174 ± 0.015 | 0.117 ± 0.010 | 0.087 ± 0.007 | 88.48 ± 12.82 | 6.64 ± 0.96 | 7.57 ± 0.92 | 0.57 ± 0.07 |
| | Min-Max | 0.211-0.332 | 0.130-0.194 | 0.090-0.134 | 0.068-0.102 | 66.27-110.69 | 4.98-8.31 | 5.98-9.17 | 0.45-0.69 |
| PFC-shoulder | M ± SEM | 0.355 ± 0.033 | 0.230 ± 0.021 | 0.166 ± 0.015 | 0.131 ± 0.012 | 159.09 ± 4.46 | 11.94 ± 0.33 | 12.52 ± 0.42 | 0.94 ± 0.03 |
| | Min-Max | 0.113-0.452 | 0.074-0.290 | 0.054-0.210 | 0.043-0.168 | 117.10-178.91 | 8.79-13.43 | 8.91-14.65 | 0.67-1.10 |
| PFC HD | M ± SEM | 0.227 ± 0.017 | 0.084 ± 0.012 | 0.026 ± 0.005 | 0.013 ± 0.003 | 23.58 ± 3.83 | 1.77 ± 0.29 | 5.46 ± 0.72 | 0.41 ± 0.05 |
| | Min-Max | 0.158-0.315 | 0.033-0.146 | 0.009-0.058 | 0.004-0.039 | 6.15-41.75 | 0.46-3.13 | 2.49-8.69 | 0.19-0.65 |
| PFC bifrontal | M ± SEM | 0.372 ± 0.050 | 0.231 ± 0.031 | 0.129 ± 0.017 | 0.075 ± 0.010 | 82.02 ± 0.83 | 6.16 ± 0.06 | 11.91 ± 0.32 | 0.89 ± 0.02 |
| | Min-Max | 0.223-0.488 | 0.137-0.302 | 0.080-0.165 | 0.047-0.096 | 80.95-86.09 | 6.08-6.46 | 10.89-12.48 | 0.82-0.94 |
| All right | M ± SEM | 0.337 ± 0.027 | 0.198 ± 0.021 | 0.120 ± 0.014 | 0.081 ± 0.011 | 91.17 ± 10.52 | 6.85 ± 0.79 | 11.22 ± 0.96 | 0.84 ± 0.07 |
| | Min-Max | 0.104-0.509 | 0.035-0.318 | 0.009-0.219 | 0.004-0.173 | 5.37-168.95 | 0.40-12.69 | 2.26-20.01 | 0.17-1.50 |

Abbreviations: $E_{MAX}$ = robust maximum electric field strength, $E_{95}$ = 95$^{th}$ percentile of electric field strength, $E_{75}$ = 75$^{th}$ percentile of electric field strength, $E_{50}$ = 50$^{th}$ percentile of electric field strength, $Foc_{75}$ = electric field volume in $cm^3$ and as percentage of total gray matter volume for 75$^{th}$ percentile of electric field strength, $Foc_{50}$ = electric field volume in $cm^3$ and as percentage of total gray matter volume for 50$^{th}$ percentile of electric field strength, HD = high definition, M = mean, PFC = prefrontal cortex, SEM = standard error of mean, SOR = supraorbital region.
